# CHIP ubiquitin ligase is involved in the nucleolar stress management

**DOI:** 10.1101/2022.05.17.492288

**Authors:** Malgorzata Piechota, Lilla Biriczova, Konrad Kowalski, Natalia A. Szulc, Wojciech Pokrzywa

**Author notes:** Correspondence should be directed to: WP, MP. These authors contributed equally to this work (Lilla Biriczova, Konrad Kowalski).

## Abstract

The nucleolus is a dynamic nuclear biomolecular condensate involved in cellular stress response. Under proteotoxic stress, the nucleolus can store damaged proteins for refolding or degradation. HSP70 chaperone is a well-documented player in the recovery process of proteins accumulated in the nucleolus after heat shock. However, little is known about the involvement of the ubiquitin-proteasome system in the turnover of its nucleolar clients. Here we show that HSP70, independently of its ATPase activity, promotes migration of the CHIP (carboxyl terminus of HSC70-interacting protein) ubiquitin ligase into the granular component of the nucleolus, specifically after heat stress. We show that while in the nucleolus, CHIP retains mobility that depends on its ubiquitination activity. Furthermore, after prolonged exposure to heat stress, CHIP self-organizes into large, intra-nucleolar droplet-like structures whose size is determined by CHIP ubiquitination capacity. Using a heat-sensitive nucleolar protein luciferase, we show that excess CHIP impairs its regeneration, probably through deregulation of HSP70. Our results demonstrate a novel role for CHIP in managing nucleolar proteostasis in response to stress.

## INTRODUCTION

The nucleolus is the largest subnuclear compartment formed by immiscible liquid phases that control ribosome biogenesis and translational capacity (Feric et al., 2016). It consists of three distinct layers: the fibrillar center (FC), dense fibrillar component (DFC) and granular component (GC), which surrounds FC and DFC, and the perinucleolar compartment (PNC) (Biggiogera et al., 1990; Feric et al., 2016). FC-DFC interface is a site where ribosomal DNA (rDNA) is actively transcribed, while DFC contains proteins such as fibrillarin, which are essential for rDNA processing (Jordan, 1984; Scheer and Hock, 1999; Smirnov et al., 2014; Lafontaine et al., 2021). GC is abundant in nucleophosmin (NPM1) and nucleolin proteins (Biggiogera et al., 1990) and acts as a site of ribosome subunits assembly (Feric et al., 2016; Kozakai et al., 2016; Mitrea et al., 2018). In addition, the nucleolus can serve as a safe harbor for proteins after exposure to environmental stimuli or stress factors. For example, labile proteins during heat stress are transported into the nucleolus, where the heat shock protein 70 (HSP70) protects them from aggregation and facilitates their extraction and refolding after stress (Nollen et al., 2001; Frottin et al., 2019). Thus, the HSP70 chaperone is essential for the maintenance of nucleolar proteostasis. Recent proteomic analysis of the nucleolus from heat-shock treated cells identified numerous proteins accumulating in nucleoli among which several belonged to the ubiquitin-proteasome system (UPS) (Azkanaz et al., 2019). UPS regulates various cellular pathways by removing unwanted and damaged proteins marked by a small protein – ubiquitin (Ub). Its attachment is mediated by the Ub-activating enzymes (E1), Ub-conjugating enzymes (E2), and Ub ligases (E3) that select target proteins. In most cases, the proteasome subsequently degrades ubiquitinated proteins (Komander, 2009; Buetow and Huang, 2016). However, little is known about the involvement of the UPS in nucleolar stress response and proteostasis maintenance. Recent studies identified numerous proteins bound to NPM1 after heat shock (Frottin et al., 2019). Their accumulation was transient, only under heat shock, and HSP70 activity was required for their dissociation from NPM1 during recovery (Frottin et al., 2019). Interestingly, several E3 ligases were detected in the aforementioned study, but further investigation of their nucleolar functions was not carried out. One of these was the quality control E3 ligase CHIP (C-terminus of Hsc70-interacting protein), the well-known HSP70 interactor. CHIP contains three tandem tetratricopeptide repeat (TPR) motifs that bind to the HSP70 and HSP90 chaperones and the catalytic U-box domain responsible for substrate ubiquitination (Ballinger et al., 1999; Jiang et al., 2001). Early work showed that heat-treated CHIP retains its ubiquitination activity and can modify substrates bound to HSP70/HSC70 (Ballinger et al., 1999; Meacham et al., 2001; Murata et al., 2001; Shimura et al., 2004; Tateishi et al., 2004; Younger et al., 2004; Stankiewicz et al., 2010). In addition, CHIP can control HSP70 levels in the client-free state through ubiquitination or by activating the transcription factor HSF1, a key regulator of the heat shock response (HSR) in eukaryotic cells (Dai et al., 2003; Qian et al., 2006). In mouse fibroblasts where CHIP was knocked out, heat-induced HSP70 activation was significantly reduced. On the other hand, its turnover rate also decreased, indicating that HSP70 and CHIP closely collaborate on degrading the chaperone’s substrates, and their interaction is also self-regulatory (Dai et al., 2003; Qian et al., 2006). However, it is unclear what is the role of CHIP while in the nucleolus and whether it also cooperates with HSP70 in maintaining nucleolar proteostasis during heat stress and recovery.

Here we show that heat shock-induced CHIP migration to nucleoli depends on HSP70 presence but not its activity. Nevertheless, functional HSP70 is essential for the release of CHIP from the nucleolus. We also noted that nucleolar CHIP could exhibit ubiquitination activity during heat stress and recovery. Specifically, CHIP is recruited to the GC compartment where it acts as a non-aggregating protein; however, its mobility becomes significantly limited when deprived of ubiquitination ability. Remarkably, CHIP localizes to specific condensates generated in the nucleolus under prolonged heat stress and whose dynamics depend on its E3 activity. To this end, we used luciferase as a stress-sensitive model protein sorted to the nucleolus during heat shock and observed that CHIP hinders its regeneration, likely in collaboration with HSP70. Our results provide the groundwork for further studies on CHIP function in a nucleolar heat stress response.

## RESULTS

### CHIP translocates to nucleoli in heat-stressed cells

To investigate the localization and function of CHIP in heat-stressed cells, we established two cell lines, based on the Flp-In system compatible HeLa and HEK-293 cells (Szczesny et al., 2018), stably overexpressing CHIP tagged with the EGFP fluorescent marker (hereafter HeLa EGFP-CHIP and HEK EGFP-CHIP cells) (Fig. S1A). In the experiment we exposed cells to heat shock at 42°C for 90 min, followed by recovery period at 37°C for 2 h (Fig. 1A). We found that EGFP-CHIP localized to nucleoli in both cell lines, specifically during heat shock, and abandoned it upon recovery (Fig.1B, Fig. S1B). None of the other tested stressors, such as arsenite, sorbitol, thapsigargin, or puromycin, induced CHIP migration into the nucleolus (Fig. S1B). We further confirmed the ability of CHIP to translocate into nucleoli in MCF7 breast cancer cells transiently transfected with EGFP-CHIP (Fig. S1C). Importantly, endogenous CHIP also accumulated in the nucleolus after heat shock in HeLa Flp-In and MCF7 cells (Fig. 1C and S1D), indicating that CHIP translocation to the nucleolus is not an artifact resulting from EGFP tagged protein overexpression. Quantification of endogeneous CHIP intensities across nuclei and nucleoli in MCF7 cells confirmed its increased nucleolar localization during heat stress (Fig. S1E).

**Fig. 1.**
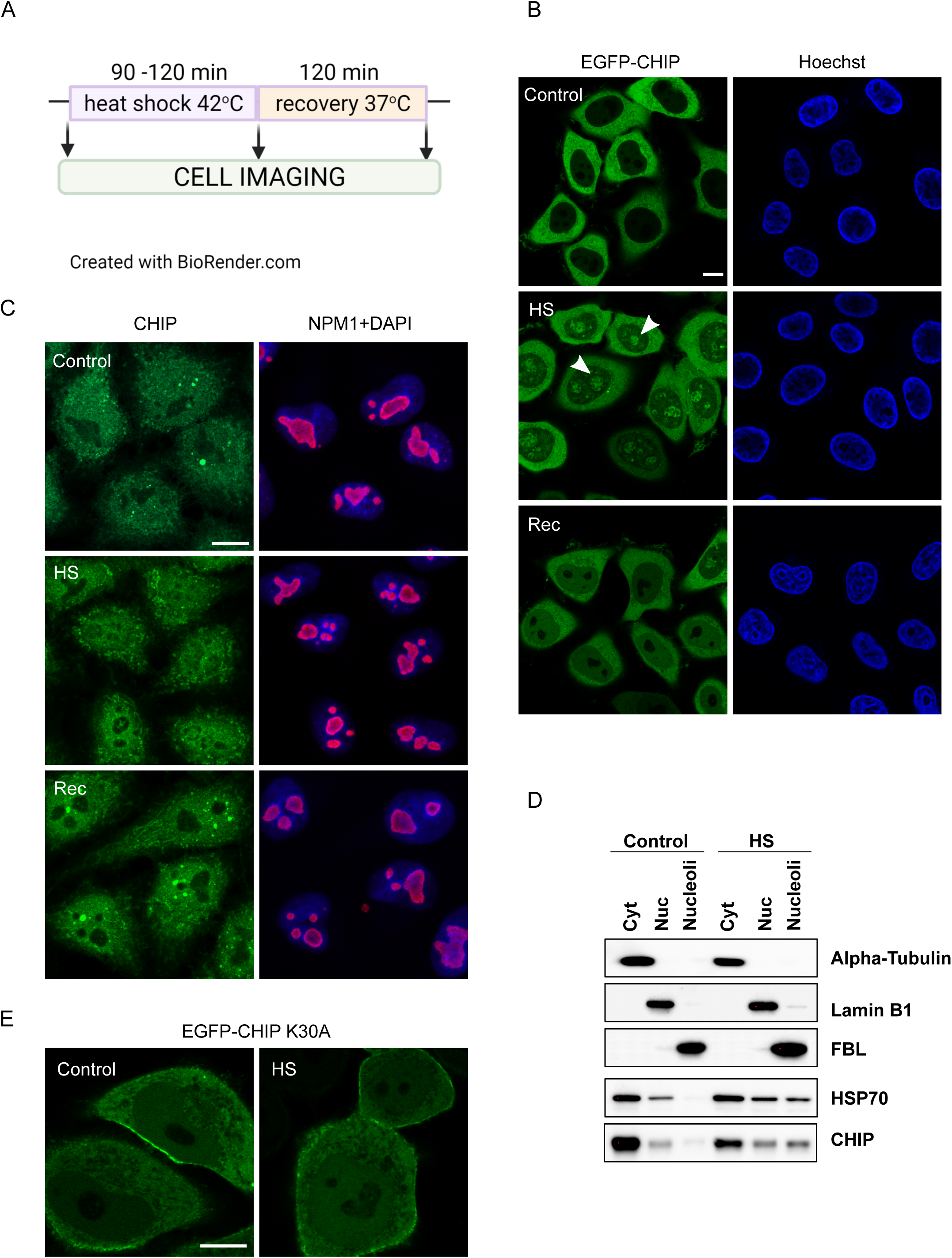
CHIP translocates to nucleoli in heat-stressed cells. A) Scheme of the heat shock assay. In most experiments, cells were exposed to heat stress at 42°C for 90 min (120 min for HEK293T cells) and transferred to 37°C for 2 h recovery. B) Overexpressed EGFP-CHIP in HeLa Flp-In cells (HeLa EGFP-CHIP cells) shows nucleolar localization after heat shock (white arrowheads). Cells were imaged live before heat shock (control), immediately after heat shock (HS), and post-heat shock recovery (Rec). Scale bar represents 10 µm. C) Endogenous CHIP can migrate to nucleoli upon heat shock. Confocal images of HeLa Flp-In cells after immunofluorescent staining for CHIP (green) and NPM1 (red) in control, heat-stressed (HS), and recovered (Rec) cells. Scale bar represents 10 µm. D) Western blot after HeLa Flp-In cell fractionation to cytoplasmic (Cyt), nucleoplasmic (Nuc), and nucleolar fractions showing CHIP and HSP70 accumulation in nucleoli after heat shock. Fractions purity was evaluated by detecting α-tubulin (cytoplasm), lamin B1 (nucleoplasm), and fibrillarin, FBL (nucleoli). E) Representative confocal images of HeLa Flp-In cells transiently expressing the EGFP-CHIP K30A show weaker translocation of this co-chaperone mutant to nucleoli upon heat shock (HS). Scale bar represents 10 µm.

Next, we fractionated HeLa Flp-In cells into cytoplasmic, nucleoplasmic, and nucleolar fractions to verify CHIP translocation into nucleoli. Consistent with our imaging data, we found elevated CHIP levels in the nucleolar fractions of heat-stressed cells (Fig. 1D). Importantly, we also detected elevated levels of HSP70 chaperone in nucleolar fractions after heat shock, which is consistent with previous reports (Pelham et al., 1984; Pelham, 1984; Welch and Feramisco, 1984; Welch and Suhan, 1986; Nollen et al., 2001; Azkanaz et al., 2019; Frottin et al., 2019). Following the suggestion that HSP70 may enter nucleoli in complex with other proteins (Frottin et al., 2019), we determined whether it modulates CHIP translocation into nucleoli. To verify this, we examined the transport of the CHIP K30A mutant, which is deficient in HSP70 binding. Indeed, HeLa Flp-In cells transiently transfected with the EGFP-CHIP K30A mutant exhibited impaired CHIP migration to nucleoli after heat shock (Fig. 1E). Therefore, we further aimed to determine the role of HSP70 in CHIP nucleolar localization.

### HSP70-dependent localization of CHIP in nucleoli

HSP70 colocalizes with CHIP in the nucleoli of heat-treated HeLa EGFP-CHIP cells, suggesting their functional cooperation (Fig. 2A). To examine the role of HSP70 in nucleolar CHIP accumulation, we lowered the HSP70 level via siRNA silencing in HeLa EGFP-CHIP cells (Fig. S2A) and applied our heat shock scheme (Fig. 1A). We observed reduced CHIP migration to the nucleoli in these cells (Fig. 2B). Furthermore, depleting HSP70 hindered CHIP exit from nucleoli during regeneration where 2 h post-heat shock more than 60% of cells still maintained CHIP in nucleoli, compared to approximately 10% of control cells (Fig. 2B and 2C).

**Fig. 2.**
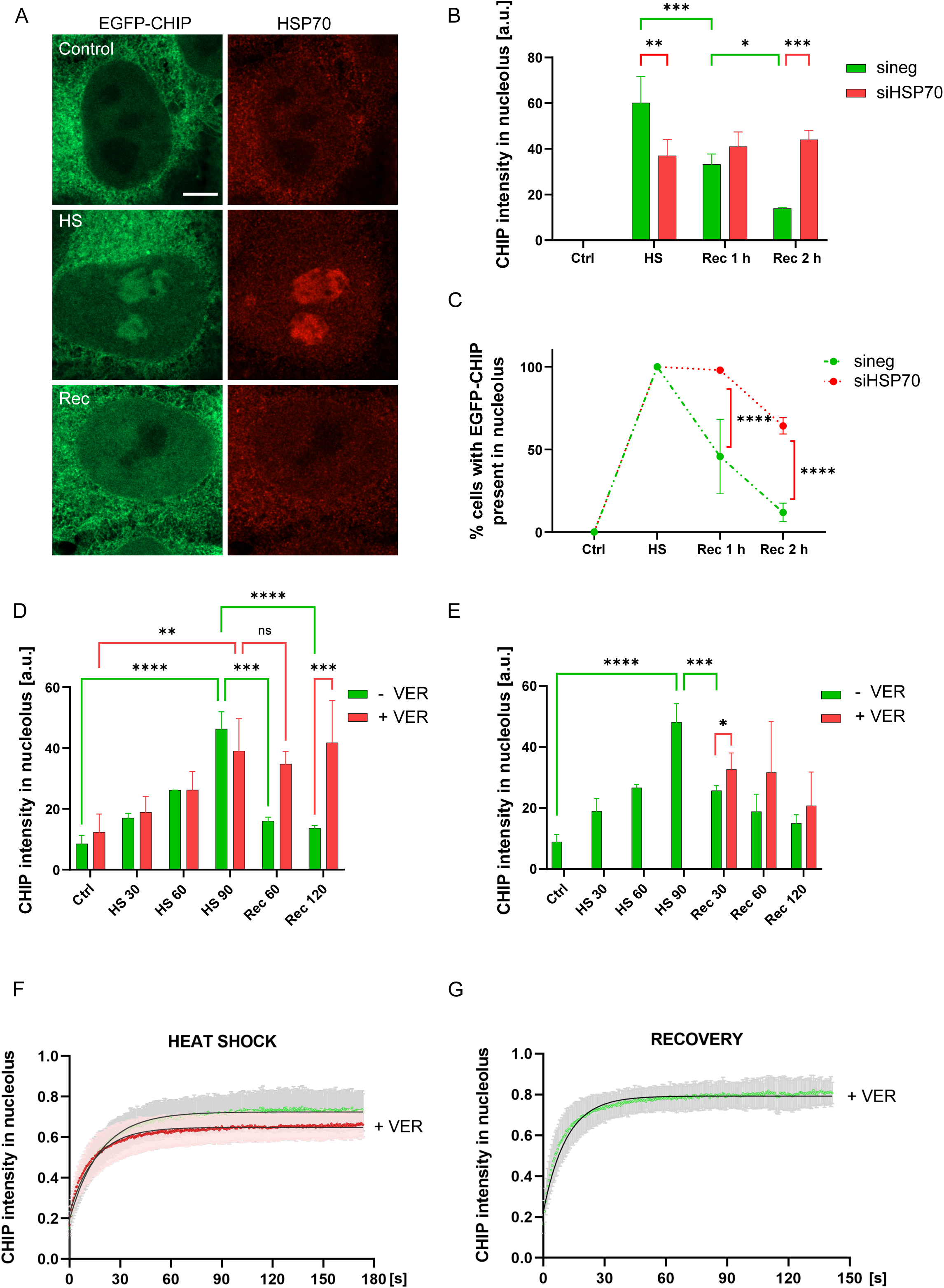
HSP70-dependent localization of CHIP in nucleoli. A) CHIP colocalizes with HSP70 upon heat shock. Confocal images (with Airyscan) of HeLa EGFP-CHIP cells after immunostaining for HSP70 (red). Scale bar represents 5 µm. B) HSP70 recruits CHIP to nucleoli during heat shock. Quantification of mean CHIP intensity in nucleoli during 90 min heat shock and 2 h-recovery in HeLa EGFP-CHIP cells upon HSP70 knockdown. Data are means of three independent experiments. Error bars show SD. Statistical significance was determined using a two-way *ANOVA* followed by Tukey’s multiple comparison test (\*\*\**P* < 0.001, \*\**P* < 0.01, \**P* < 0.05). C) Quantification of the percentage of cells with CHIP present in nucleoli during heat shock and 2 h-recovery in HeLa EGFP-CHIP cells upon HSP70 knockdown. Data are means of three independent experiments. Error bars show SD. Statistical significance was determined using a two-way *ANOVA* followed by Tukey’s multiple comparison test (\*\*\*\**P* < 0.0001). D) HSP70 inhibition by VER does not affect CHIP migration to nucleoli during heat shock but blocks its release during recovery. HeLa EGFP-CHIP cells were treated with 40 µM VER before heat shock, and CHIP intensity was measured in nucleoli in control cells during heat shock and 2 h-recovery. Data are means of three independent experiments. Error bars show SD. Statistical significance was determined using a two-way *ANOVA* followed by Tukey’s multiple comparison test (\*\*\*\**P* < 0.0001, \*\*\**P* < 0.001, \*\**P* < 0.01, *ns P* > 0.05). E) HSP70 inhibition by VER during post-heat shock recovery only slightly affects CHIP clearance from nucleoli. HeLa EGFP-CHIP cells were exposed to 90 min heat shock and treated with 40 µM VER before transferring them for the 2 h-recovery. CHIP intensity was measured in nucleoli in control cells during heat shock and recovery. Data are means of four independent experiments. Error bars show SD. Statistical significance was determined using two-tailed unpaired t-tests for pairwise comparisons (\*\*\*\**P* < 0.0001, \*\*\**P* < 0.001, \**P* < 0.05). F) CHIP maintains high mobility in the nucleolus upon heat shock. Analysis of FRAP kinetics of EGFP-CHIP in the nucleolus of untreated (green) or treated with 40 µM VER (red) HeLa EGFP-CHIP cells during heat shock. Points show mean values from 9 or 19 nucleoli analysis from untreated or VER-treated cells, respectively. Error bars show SD (grey for untreated cells, pink for VER-treated cells). Fitting curves are shown in black. G) CHIP dynamics quantification in HeLa EGFP-CHIP cells during post-heat shock recovery in the presence of 40 µM VER. FRAP kinetics were measured in 15 nucleoli after 1 h-recovery. Error bars show SD. A fitting curve is shown in black.

We next monitored CHIP localization during heat shock and recovery in the presence of VER-155008 (hereafter VER), a small molecule inhibitor of HSP70, to test whether HSP70 activity is required for CHIP translocation to nucleoli. Notably, we used 40 µM VER in HeLa EGFP-CHIP and MCF7 cells as this concentration was successfully applied to inhibit HSP70 activity in HeLa cells during recovery from the 3 h heat shock (Mediani et al., 2019). We observed CHIP levels gradually increasing in the nucleoli of HeLa EGFP-CHIP and MCF7 cells during heat shock, implying that its nucleolar migration was not affected by HSP70 inhibition (Fig. 2D and S1E). However, continuous VER treatment during heat shock and recovery blocked CHIP release during recovery, which resembled the effect of HSP70 depletion (Fig. 2D and S1E). These results suggest that HSP70 recruits CHIP in an activity-independent manner upon entry to the nucleolus, but its functional operability in this compartment is required for the recovery process and consequent CHIP release. When VER was provided only during the recovery stage, CHIP clearance from nucleoli was only slightly reduced (Fig. 2E), indicating that CHIP trapping in nucleoli depends primarily on the functionality of the HSP70 during heat shock.

Therefore, we wanted to determine whether CHIP in the nucleolus acts as a functional protein or as the HSP70 substrate, misfolded upon heat shock. Based on observations that misfolded proteins acquire low mobility in the nucleolus (Azkanaz et al., 2019; Frottin et al., 2019), we analyzed CHIP nucleolar fraction mobility in untreated cells and in the presence of VER by recording fluorescence recovery after photobleaching (FRAP). Approximately 70% of EGFP-CHIP sequestered in nucleoli after heat shock was mobile, and HSP70 inhibition did not significantly reduce its dynamics (Fig. 2F). CHIP mobility was also unchanged after 1 h post-heat shock recovery in the presence of VER (Fig. 2G). This indicates that CHIP can form a functional, unaggregated protein in the nucleoli.

### Nucleolar CHIP colocalizes with the NPM1-containing granular component (GC) phase

The organization of the nucleolus, involving all three layers, is essential for its role in ribosome biogenesis (Huang, 2002; Krüger et al., 2007; Riback et al., 2020). In turn, the GC is thought to be the main phase supporting misfolded proteins translocated there during proteotoxic stress (Azkanaz et al., 2019; Frottin et al., 2019). To study CHIP specific localization in the nucleolus, we performed colocalization analysis using confocal microscopy with the Airyscan detector, contributing to the improved image resolution (Wu and Hammer, 2021). HeLa EGFP-CHIP cells were stained for NPM1, a GC marker, and fibrillarin (FBL), a DFC marker. Under heat shock and recovery conditions, with or without VER, CHIP colocalized with NPM1 (Fig. 3A and 3C) and, to a much lesser extent, with FBL (Fig. 3B and 3C), suggesting potential CHIP involvement in protein quality control processes in GC.

**Fig. 3.**
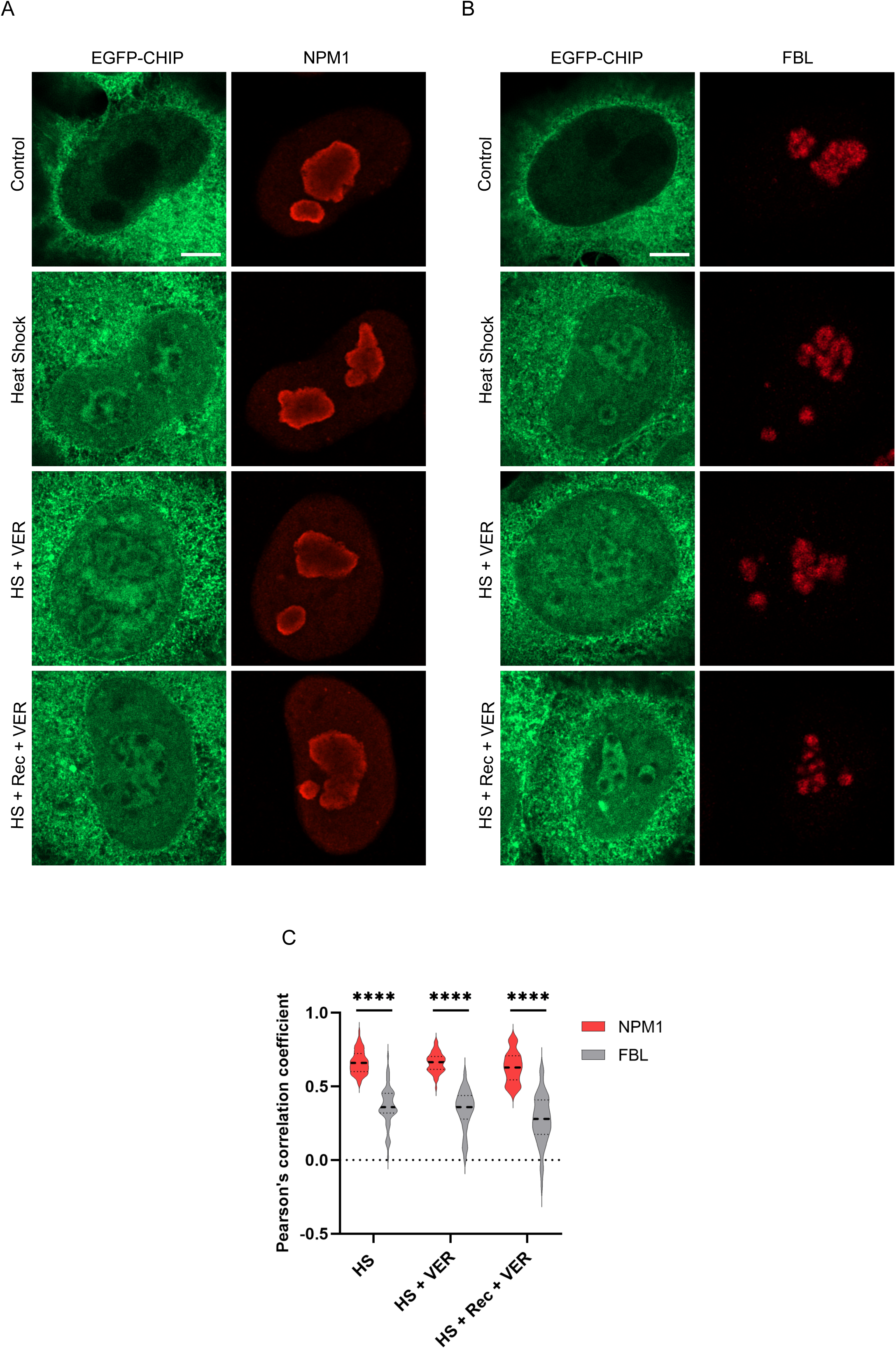
Nucleolar CHIP colocalizes with the NPM1-containing granular component (GC) phase. A) Confocal images (with Airyscan) of HeLa EGFP-CHIP cells immunostained for NPM1. Cells were exposed to the following conditions: heat shock, heat shock in the presence of 40 µM VER, or treated with VER throughout heat shock and recovery (HS + Rec + VER), followed by immunostaining. Scale bar represents 5 µm. B) Confocal images (with Airyscan) of HeLa EGFP-CHIP cells immunostained for FBL. Cells were exposed to the following conditions: heat shock, heat shock in the presence of 40 µM VER, or treated with VER throughout heat shock and recovery (HS + Rec+ VER), followed by immunostaining. Scale bar represents 5 µm. C) Quantification of the degree of colocalization of EGFP-CHIP and NPM1 and EGFP-CHIP and FBL using Pearson’s correlation coefficient. Violin plots show the data from 31 to 70 nucleoli analyzed per condition. Two-tailed unpaired t-tests were used for comparisons. Statistical significance level \*\*\*\**P* < 0.0001.

### CHIP import to nucleoli is not induced by nucleolar stress *per se*

To investigate whether CHIP migration to the nucleolus can be triggered as a result of nucleolus impairment, we treated cells with low doses of the transcription inhibitor Actinomycin D (hereafter Act D), which alters the distribution of multiple nucleolar proteins, resulting in the formation of nucleolar caps (Reynolds et al., 1964; Shav-Tal et al., 2005). Act D altered the morphology of nucleoli, causing their circularization, reduction in size, and the formation of FBL nucleolar caps (Fig. S3A and S3B). However, it did not induce CHIP migration to nucleoli (Fig. 4A-C). These results support the concept that CHIP is involved in the nucleolar heat stress response process rather than, for example, suppressing rRNA transcription defects. While treatment with Act D prior to heat shock did not affect CHIP migration to nucleoli (Fig. 4B and 4C), it altered CHIP distribution, which more prominently overlapped with Act D-induced NPM1 ring formations (Fig. 4D). In addition, in cells exposed to Act D, CHIP exit from the nucleolus during the 2 h heat stress recovery was partially impaired (Fig. 4B, 4C and S3C). This observation suggests that proper nucleolar assembly may be necessary for CHIP dynamics.

**Fig. 4.**
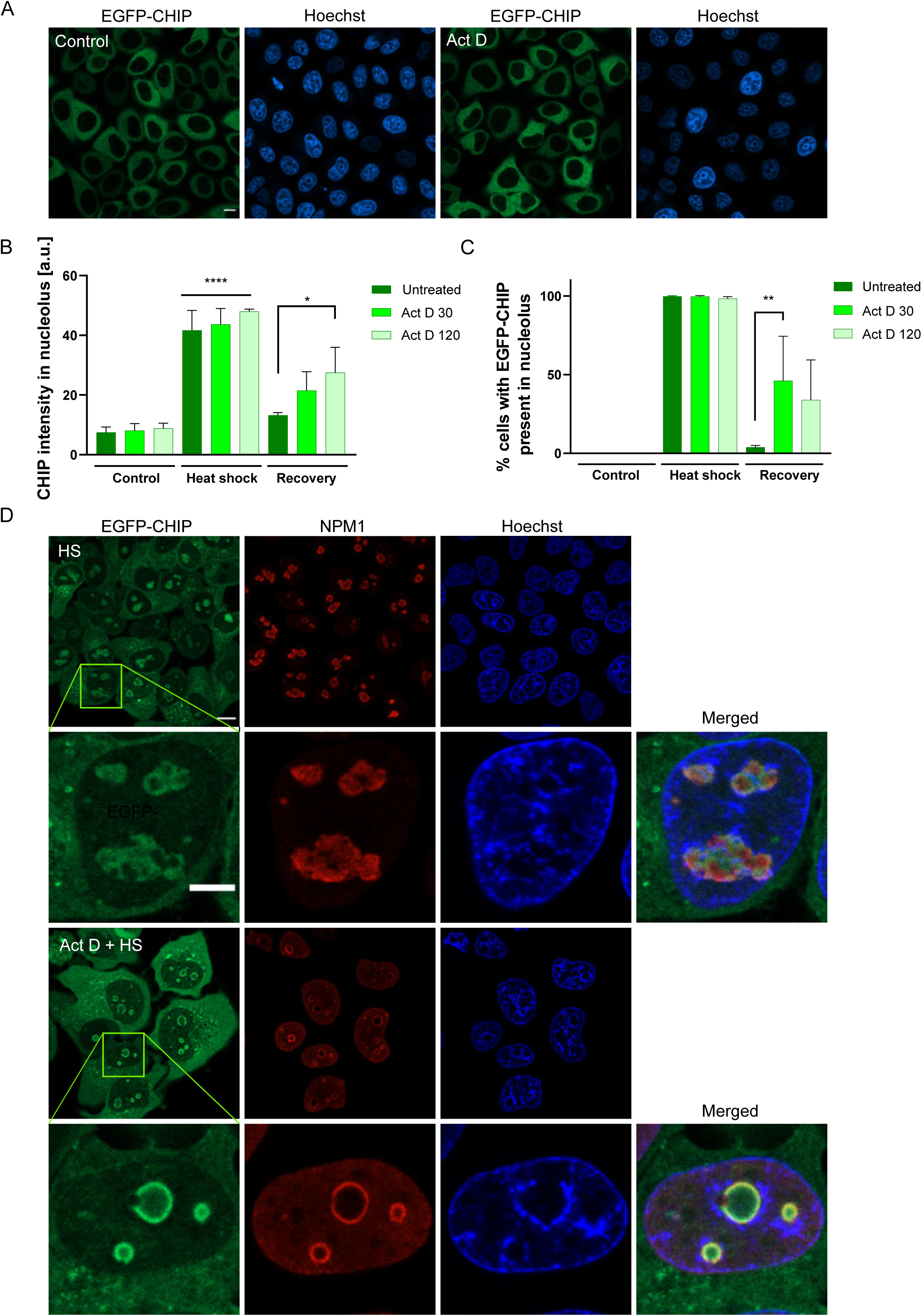
CHIP import to nucleoli is not induced by nucleolar stress *per se*. A) Actinomycin D (Act D) does not induce CHIP migration to nucleoli in HeLa EGFP-CHIP cells. Cells were treated with 0.05 µg/ml Act D for 30 min or 2 h and imaged live. Representative confocal images of cells after 2 h Act D treatment. Scale bar represents 10 µm. B) Pretreatment with 0.05 µg/ml Act D before heat shock does not affect CHIP migration to nucleoli during heat shock but impairs its exit. Quantification of mean CHIP intensity in nucleoli after 90 min heat shock and 2 h-recovery in HeLa EGFP-CHIP cells pretreated with Act D for 30 min or 2 h. Data are means of three independent experiments. Error bars show SD. Statistical significance was determined using a one-way *ANOVA* followed by Dunnett’s multiple comparison tests (\*\*\*\**P* < 0.0001, \**P* < 0.05). C) Pretreatment with 0.05 µg/ml Act D before heat shock does not affect CHIP migration to nucleoli during heat shock but impairs its exit. Quantification of the percentage of cells with CHIP present in nucleoli after 90 min heat shock and 2 h-recovery in HeLa EGFP-CHIP cells pretreated with Act D for 30 min or 2 h. Data are means of three independent experiments. Error bars show SD. Statistical significance was determined using a one-way *ANOVA* followed by Dunnett’s multiple comparison tests (\*\**P* < 0.01). D) Treatment with Act D prior to heat shock alters CHIP distribution in nucleoli. HeLa EGFP-CHIP cells were pretreated with 0.05 µg/ml Act D for 2 h before heat shock, followed by immunostaining for NPM1 and confocal imaging. Representative images and their magnified views of cells after heat shock (HS) *vs*. cells treated with Act D before heat shock (Act D + HS) are shown.Scale bars represent 10 µm or 5 µm (magnified views).

### CHIP activity promotes its dynamics in the nucleolus

We used a modified *in vitro* ubiquitination assay using total cell lysate as a CHIP source to verify if CHIP activity is maintained in nucleoli. This assay is based on the ability of an E3 ligase to self-ubiquitinate in the presence of the complete ubiquitination enzymatic cascade, namely E1 ubiquitin-activating enzyme, E2 ubiquitin-conjugating enzyme, and E3 ligase of interest, with the addition of ubiquitin and ATP, and was repeatedly used to assess CHIP activity in other studies (Murata et al., 2001; Das et al., 2021). We found that neither heat shock nor recovery period affected CHIP ubiquitination activity (Fig. 5A). This is in line with the mobile and unaggregated nucleolar fraction of CHIP (Fig. 2F and G) and implies its capability of performing self- or substrates’ ubiquitination. We also investigated whether CHIP activity is required for its translocation using the catalytically-inactive CHIP H260Q mutant (Hatakeyama et al., 2001). We found that the activity of CHIP is not indispensable for heat shock-induced migration to the nucleolus (Fig. 5B). However, FRAP analysis of the nucleolar CHIP H260Q mutant showed a decrease in its dynamics compared to CHIP WT, suggesting that its propensity to aggregate is likely mediated by the loss of ubiquitination activity (Fig. 5C and 5D).

**Fig. 5.**
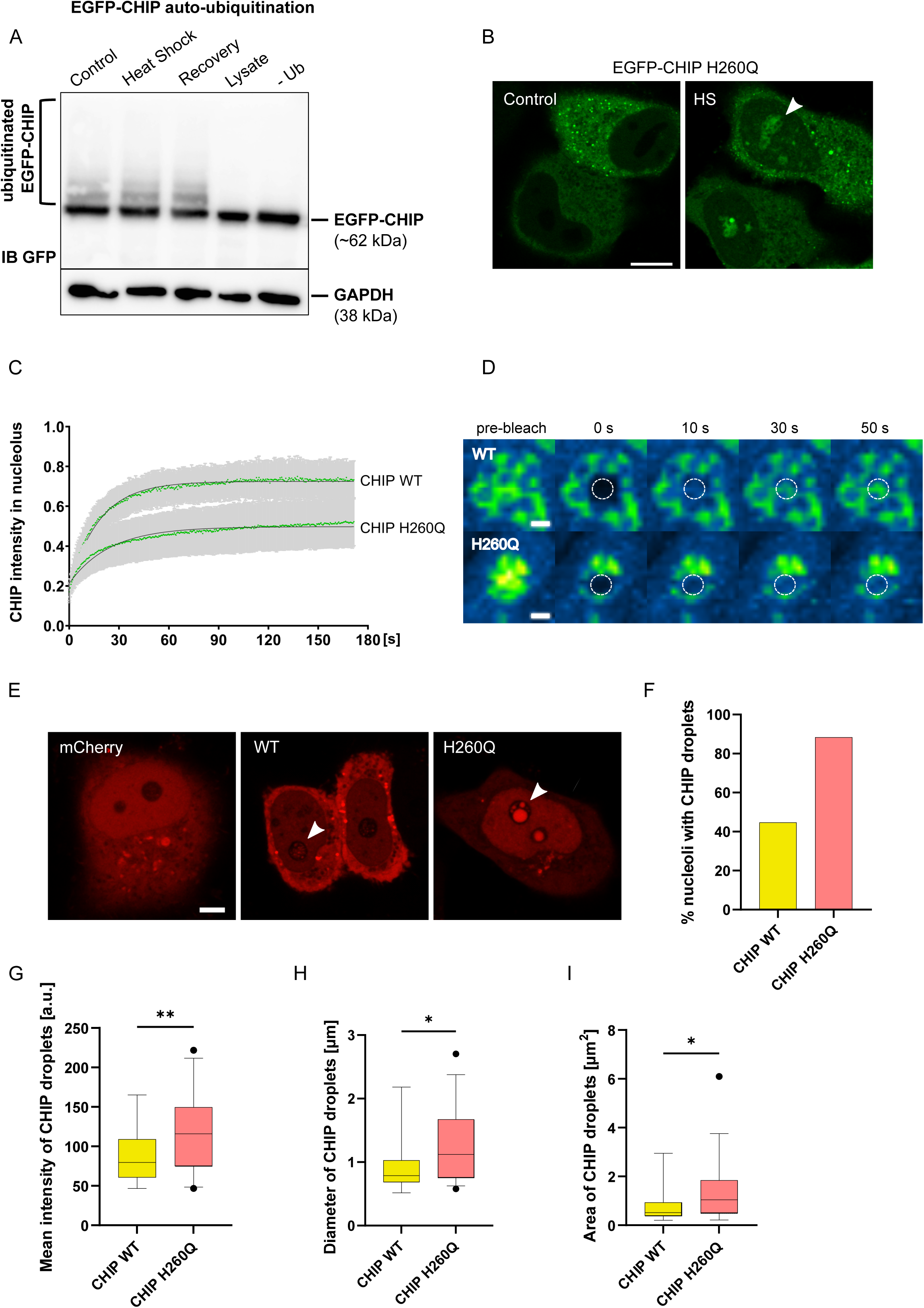
CHIP activity promotes its dynamics in the nucleolus. A) Heat shock and post-heat shock recovery do not affect CHIP ubiquitination activity. Western blot depicting CHIP auto-ubiquitination following *in vitro* ubiquitination assay. HEK EGFP-CHIP cells were exposed to 90 min heat shock and 1 h-recovery. After treatment, cell lysates were used for *in vitro* ubiquitination assay. The assays performed in the presence of the lysate without other components in the reaction mixture (lysate) or without the addition of ubiquitin (-Ub) were served as negative controls. Protein samples were resolved via SDS-PAGE and immunoblotted with anti-GFP and GAPDH (loading control) antibodies. B) Inactive CHIP H260Q mutant can migrate to nucleoli during heat shock. Representative confocal images of HeLa Flp-In cells transiently expressing the EGFP-CHIP H260Q mutant in control conditions and after heat shock (HS). The arrowhead points at the nucleolus containing mutant CHIP. Scale bar represents 10 µm. C) The CHIP H260Q mutant shows reduced mobility in nucleoli. FRAP kinetics of CHIP H260Q compared to CHIP WT in nucleoli of heat-shocked cells. The mutant CHIP mobility was measured in HeLa Flp-In cells after transient expression of EGFP-CHIP H260Q. FRAP analysis of EGFP-CHIP WT was originally shown in Fig. 2F and is displayed here again for comparison with CHIP H260Q recovery curves. CHIP H260Q FRAP kinetics were measured in 12 nucleoli. Error bars show SD. Fitting curves are shown in black. D) Confocal images of nucleolar EGFP-CHIP WT (upper panel) and EGFP-CHIP H260Q mutant (bottom panel), showing movie frames before bleaching and 0, 10, 30, and 50 s after bleaching in the FRAP assays. The bleached region of interest is marked with circles. Differences in CHIP intensities are displayed in pseudo-colored images using Green Fire Blue LUT (look-up table) in ImageJ software. Scale bars represent 2 µm. E) Large droplet-like structures are preferentially formed by the CHIP H260Q mutant in nucleoli of cells exposed to prolonged heat shock. Representative confocal images of HeLa Flp-In cells transiently expressing mCherry, mCherry-CHIP WT, and mCherry-CHIP H260Q mutant and treated with overnight heat shock. Arrowheads point at CHIP intra-nucleolar assemblies. Scale bar represents 5 µm. F) Quantification of the percentage of nucleoli with CHIP droplet-like structures after overnight heat shock in cells transiently expressing CHIP WT or the CHIP H260Q mutant. Data were collected from two independent experiments: 38 and 43 nucleoli with CHIP WT and H260Q mutant, respectively. G) Boxplot of the mean intensities of CHIP droplet-like structures inside nucleoli of cells transiently expressing CHIP WT or the CHIP H260Q mutant after overnight heat shock. Note that if there were several droplets inside nucleoli, the intensity was measured for the brightest one. 17 CHIP WT and 38 CHIP H260Q droplets across two biological repeats were analyzed. The line in the middle of the box is plotted at the median. Whiskers extend from the 5^th^ to 95^th^ percentiles. Two-tailed Welch’s t-test was used for comparison, \*\**P* < 0.01. H) Quantification of the diameters of CHIP droplet-like structures inside nucleoli of cells transiently expressing CHIP WT or the CHIP H260Q mutant after overnight heat shock. Note that if there were several droplets inside nucleoli, the measurement was performed for the biggest one. 17 CHIP WT and 38 CHIP H260Q droplets across two biological repeats were analyzed. The line in the middle of the box is plotted at the median. Whiskers extend from the 5^th^ to 95^th^ percentiles. For comparison two-tailed Mann-Whitney test was used, \**P* < 0.05. I) Boxplot of the areas of CHIP droplet-like structures inside nucleoli of cells transiently expressing CHIP WT or the CHIP H260Q mutant after overnight heat shock. Note that if there were several droplets inside nucleoli, the measurement was performed for the biggest one. 17 CHIP WT and 38 CHIP H260Q droplets across two biological repeats were analyzed. The line in the middle of the box is plotted at the median. Whiskers extend from the 5^th^ to 95^th^ percentiles. For comparison two-tailed Mann-Whitney test was used, \**P* < 0.05.

Nucleoli are sites for immobilization of proteins under heat stress, leading to occurrence of nucleolar foci with an amyloid-like character (Wang et al., 2019). To gain better insight into the long-term impact of proteotoxic stress on CHIP association with nucleoli and the consequences of its inactivity on this process, we subjected cells to prolonged heat shock. Interestingly, sizeable intra-nucleolar CHIP droplet-like structures could be observed after overnight heat shock in cells expressing the CHIP H260Q mutant, outnumbering their WT protein counterparts (Fig. 5E-I). These differences between CHIP WT and mutant assemblies may stem from the alterations in CHIP H260Q dynamics within the nucleolus (Fig. 5C and D). However, we do not know specific biophysical state and function of these structures.

### CHIP overexpression affects the nucleolar luciferase recovery

To investigate whether CHIP abundance in nucleoli can affect the fate of misfolded proteins sorted there, we employed thermolabile luciferase as a model protein since early reports showed that CHIP could control its heat shock-denatured state (Ballinger et al., 1999; Murata et al., 2001; Kampinga et al., 2003; Rosser et al., 2007). To this end, we used the HEK293T cell line permanently expressing a fusion protein of firefly luciferase and heat-stable green fluorescent protein (GFP) carrying an N-terminal nuclear localization signal (hereafter luciferase) (Frottin et al., 2019). This luciferase translocates to nucleoli after heat shock and relocates to the nucleoplasm during recovery. We verified a similar luciferase shuttle using our heat shock/recovery scheme (Fig. 1A) and noted that transiently overexpressed CHIP (tagged with mCherry) colocalizes with luciferase during heat shock (Fig. 6A). To investigate the role of CHIP in nucleolar luciferase processing, we expressed its K30A and H260Q mutants, which inhibit HSP70 binding or CHIP activity, respectively, in the aforementioned HEK293T cell line. As a proxy for luciferase abundance and regeneration, we analyzed the number of its foci in nucleoli during heat shock and the 6 h recovery period (Fig. S4A). Luciferase foci number decreased progressively during the recovery, but in cells expressing specifically CHIP WT or CHIP H260Q, their regeneration was slower than in untransfected and mCherry controls (Fig. S4A). Notably, in cells expressing the CHIP H260Q mutant luciferase recovery was not completed within the experimental 6 h time frame. This could be due to the high number of cells containing heat shock-induced luciferase foci and their presence in about 20% of non-heat shocked cells, suggesting that loss of CHIP activity had a potent destabilizing impact on luciferase. Therefore, we decided to normalize our data to correct for the differences in the number of luciferase foci during heat shock and control conditions, focusing explicitly on the ability of CHIP variants to affect luciferase nucleolar regeneration. Our analysis revealed that the elevated CHIP level induced a delay in the dissolution of nucleolar luciferase foci during recovery (Fig. 6B). Overexpression of CHIP WT and CHIP H260Q had the most potent effect on reducing luciferase exit from the nucleolus, and there was no difference in the rate of luciferase recovery between the two variants. In contrast, overexpression of the CHIP K30A mutant exerted a marginal effect on this process (Fig. 6B). When we transfected cells with lower amounts of plasmids to induce milder overexpression of CHIP variants and examined the first two hours of recovery from heat shock, we still observed comparable inhibition of decline of luciferase foci during recovery by CHIP WT and the H260Q mutant and no significant effect of CHIP K30A (Fig. S4B). We assumed that this was due to the inefficient transport of CHIP K30A to nucleoli, as in HeLa Flp-In cells (Fig. 1E). Surprisingly, we found comparable redistribution of all CHIP variants to nucleoli during heat shock, suggesting an alternative pathway for CHIP recruitment to nucleoli unaccompanied by HSP70 in HEK293T cells (Fig. 6C). Hence, the above results suggest that the slowed resolution of luciferase foci in nucleoli may be related to cross-talk between CHIP and HSP70.

**Fig. 6.**
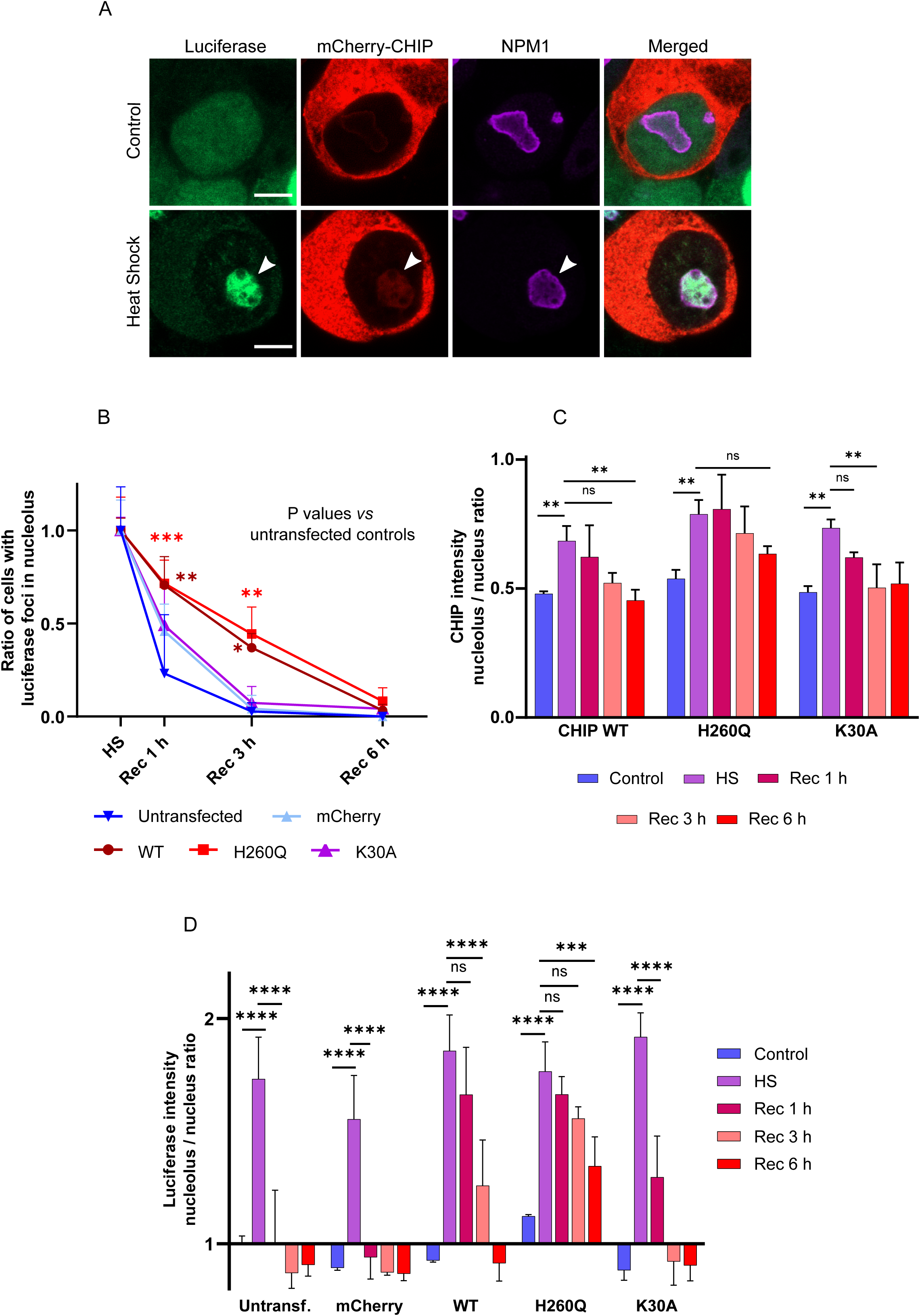
CHIP overexpression affects the nucleolar luciferase recovery. A) Nucleolar CHIP colocalizes with luciferase after heat shock. HEK293T cells stably expressing luciferase were transfected with mCherry-CHIP and subject to heat shock for 2 h. After treatment cells were immunostained for NPM1 and imaged using the Airyscanning technique. Arrowheads show overlapped signals of luciferase (green), mCherry-CHIP (red) and NPM1 (purple). Scale bars represent 5 µm. B) Luciferase foci dissolution during post-heat shock recovery is slower in cells expressing CHIP WT or CHIP H260Q but not in cells expressing CHIP K30A. HEK293T cells permanently expressing luciferase were transfected with vectors encoding for mCherry, mCherry-CHIP WT, mCherry-CHIP H260Q and mCherry-CHIP K30A. 24 h after transfection cells were subject to 2 h heat shock and the recovery was monitored for 6 h afterward. Cells were imaged live using confocal microscopy. Luciferase foci were counted in untransfected and mCherry-expressing cells (control groups) and cells expressing the appropriate CHIP variant. The percentage of cells with nucleolar luciferase foci was determined for each condition. Data were normalized to correct for cell percentage differences after heat shock and in control conditions between experimental groups and are expressed as the ratio of % cells with luciferase foci at the specific time point to % cells with luciferase foci upon heat shock calculated for a given group. Data are means of three independent experiments. Error bars represent SD. For statistical comparison a two-way *ANOVA* with post hoc Tukey’s test was used (\*\*\**P* < 0.001, \*\**P* < 0.01, \**P* < 0.05). C) CHIP is redistributed to nucleoli during heat shock and leaves this compartment during recovery. During recovery, the CHIP H260Q mutant’s exit from nucleoli is the slowest compared to CHIP WT and CHIP K30A. HEK293T cells permanently expressing luciferase were transfected with vectors encoding for mCherry-CHIP WT, mCherry-CHIP H260Q and mCherry-CHIP K30A and treated with 2 h heat shock followed by 6 h recovery. Cells during treatments were imaged live using confocal microscopy. Images were analyzed for the mean mCherry intensities as a proxy for CHIP concentrations in the nucleoli and nuclei, and the relative intensities were quantified. Data are means of three independent experiments. Error bars represent SD. For statistical comparison a two-way *ANOVA* with post hoc Tukey’s test was used (\*\**P* < 0.01, *ns P* > 0.05). D) CHIP WT and CHIP H260Q overexpression disrupt the regeneration rate of luciferase in nucleoli. HEK293T cells permanently expressing luciferase were transfected with vectors encoding for mCherry, mCherry-CHIP WT, mCherry-CHIP H260Q, and mCherry-CHIP K30A and treated with 2 h heat shock followed by 6 h recovery. Confocal images of live cells were taken after indicated time points and images were analyzed for mean intensities of GFP-tagged luciferase in whole nucleoli relative to nuclei. Data are means of three independent experiments. Error bars represent SD. For statistical comparison a two-way *ANOVA* with post hoc Tukey’s test was used (\*\*\*\**P* < 0.0001, \*\*\**P* < 0.001, *ns P* > 0.05).

We next assessed luciferase levels as the ratio of its intensity between nucleolus and nucleoplasm, as measured immediately after heat shock and during the 6 h recovery period. The distribution of luciferase in untransfected or mCherry-transfected control cells was predominant in the nucleoplasm already at the initial stage of recovery. However, in cells overexpressing CHIP WT, the nucleolar luciferase signal was still noticeable after 3 h of recovery, again indicating that the regeneration rate of luciferase was disrupted (Fig. 6D). While the CHIP K30A mutant showed the least disruption in the redistribution of luciferase, the CHIP H260Q mutant resulted in its most extended nucleolar persistence (Fig. 6D). We also observed that CHIP was leaving the nucleoli during recovery, concomitantly with nucleolar luciferase disappearance, with the slowest rate for the CHIP H260Q mutant (Fig. 6C). Thus, we assume that CHIP-dependent ubiquitination may contribute to luciferase processing in nucleoli and regeneration efficiency.

As prolonged heat shock was shown to compromise nucleolar quality control and inhibit luciferase regeneration (Frottin et al., 2019), we set out to investigate the effects of CHIP on regeneration under these conditions. We measured luciferase intensity in nuclei and nucleoli and monitored the number of luciferase foci during the 6 h heat shock at 42°C. Control cells and cells expressing CHIP K30A, but not cells expressing CHIP WT and H260Q, were capable of almost complete dissolution of luciferase foci (Fig. S5A, S5B). However, we observed sustained sequestration of CHIP H260Q into nucleoli after prolonged heat stress (Fig. S5C). Thus, we concluded that CHIP repressed rather than enhanced nucleolar luciferase regeneration. Furthermore, our results on CHIP K30A suggest that the interaction of CHIP with HSP70 may play a role in modulating the nucleolus regeneration capacity and CHIP translocation to the nucleoplasm.

### HSP70 inhibition aggravates the negative effect of CHIP on luciferase regeneration

We next examined the effect of CHIP on luciferase regeneration in the presence of VER, the HSP70 inhibitor. Cells were treated with VER only during post-heat shock recovery, and the number of nucleolar luciferase foci was measured after 1 h and 2 h of recovery. In untransfected cells, we did not record any impact of VER on luciferase regeneration. Cells overexpressing CHIP WT showed mildly impaired nucleolar luciferase regeneration in the presence of VER, which became apparent after the second hour of recovery compared to condition where it was absent (Fig. 7A). However, overexpression of the CHIP K30A in cells with added VER had a more disruptive effect relative to untreated cells (Fig. 7B). The result for the CHIP K30A mutant was unexpected as, unlike the WT protein, it should not interfere with the HSP70 function, which may suggest the emergence of additional effects associated with the chaperone inhibition. Since the negative effect on luciferase regeneration in CHIP H260Q-expressing cells was also potently enhanced by HSP70 inhibition, we speculate that protection against protein aggregation in the nucleolus requires a balance between HSP70 and E3 CHIP activity.

**Fig. 7.**
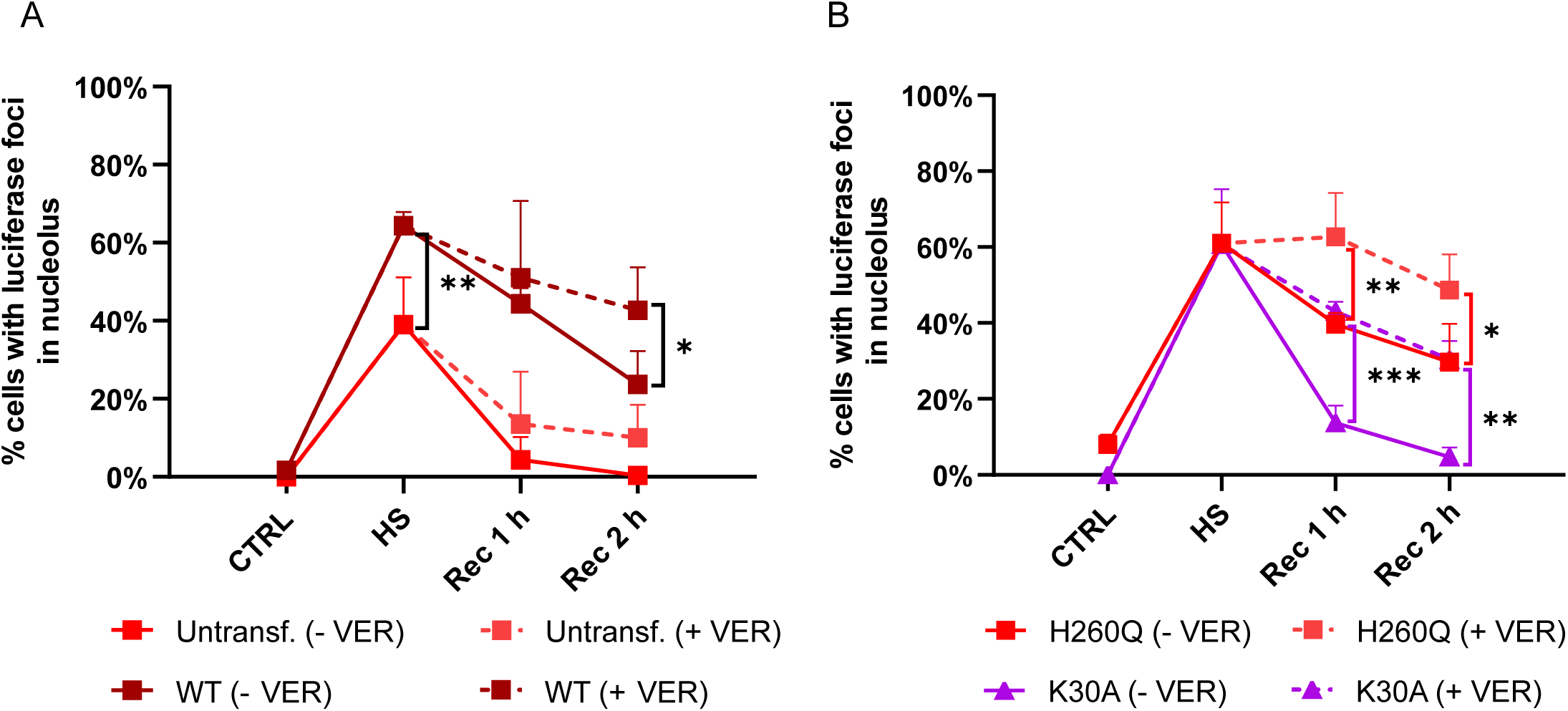
HSP70 inhibition aggravates the negative effect of CHIP on luciferase regeneration. A) Cells overexpressing CHIP WT show mildly impaired nucleolar luciferase regeneration in the presence of VER. HEK293T cells stably expressing luciferase were transfected with the vectors encoding for mCherry-CHIP variants. 24 h after transfection cells were subject to 2 h heat shock and recovery. Prior to the recovery period cells were treated with 40 µM VER or recovery was initiated without the compound treatment. The plot shows the quantification of nucleolar luciferase foci in untransfected cells and cells expressing CHIP WT imaged by confocal microscopy. Data are means of three independent experiments and are expressed as a % of total cell counts. Error bars represent SD. For statistical comparison, a two-way *ANOVA* with post hoc Tukey’s test was used (\*\**P* < 0.01, \**P* < 0.05). B) Overexpression of the CHIP K30A or CHIP H260Q mutants cause more disruptive effects on luciferase regeneration in the presence of VER. HEK293T cells stably expressing luciferase were transfected with the vectors encoding for mCherry-CHIP variants. Prior to the recovery period, cells were treated with 40 µM VER or recovery was initiated without the compound treatment. The plot shows the quantification of nucleolar luciferase foci in cells expressing CHIP K30A and CHIP H260Q imaged by confocal microscopy. Data are means of three independent experiments and are expressed as a % of total cell counts. Error bars represent SD. For statistical comparison a two-way *ANOVA* followed by Tukey’s multiple comparison test was used (\*\*\**P* < 0.001, \*\**P* < 0.01, \**P* < 0.05).

## DISCUSSION

The nucleolus possesses numerous functions, including ribosome biogenesis, nuclear organization, regulation of global gene expression, and energy metabolism (Cerqueira and Lemos, 2019). It also responds to multiple stresses, such as hypoxia, pH fluctuations, redox stress, DNA damage, or proteasome inhibition, and acts as a protein quality control center that can mitigate heat shock-induced proteotoxicity (Mekhail et al., 2005; Latonen et al., 2011; Audas et al., 2012; Yang et al., 2016; Lindström et al., 2018; Alberti and Carra, 2019; Azkanaz et al., 2019; Frottin et al., 2019; Mediani et al., 2019; Szaflarski et al., 2022). In our studies, we focused on the latter function. It was previously suggested that the nucleolus creates a favorable environment for the HSP70-mediated protection and recovery of heat stress-sensitive proteins (Nollen et al., 2001; Azkanaz et al., 2019; Frottin et al., 2019; Mediani et al., 2019). These include the epigenetic modifier family of Polycomb group (PcG) proteins and the exogenous thermolabile luciferase (Azkanaz et al., 2019; Frottin et al., 2019). In addition, Frottin et al. demonstrated reversible accumulation of CDK1 and BRD2 proteins in the nucleolus under heat stress, and Mediani et al. pointed to DRiPs (defective ribosomal products) accumulating in the nucleolus that undergoes reversible amyloidogenesis after heat shock or proteasome inhibition (Frottin et al., 2019; Mediani et al., 2019). To better understand the proteotoxic stress-dependent management of proteins in the nucleolus, we set out to study the protein quality control ubiquitin ligase CHIP, which is well known for its role in ubiquitination of HSP70 substrates, and whose presence in the nucleolus after heat stress has been reported in recent proteomic analyses (Demand et al., 2001; Petrucelli et al., 2004; Joshi et al., 2016; Azkanaz et al., 2019; Frottin et al., 2019).

We found that CHIP translocation to the nucleolus was caused by heat stress but not by Act D, ruling out a direct CHIP response to transcriptional stress and inhibition of rRNA transcription. CHIP migration was partially dependent on HSP70; however, its chaperone activity was not required. Of note, it was previously reported that the HSP70 inhibitor, VER, does not inhibit heat shock-induced nucleolar accumulation of the HSP70 substrate, PcG protein (GFP::CBX2) (Azkanaz et al., 2019). VER competes with ATP and ADP for binding to HSP70 and reduces the rate of nucleotide association and ATP-induced substrate release (Schlecht et al., 2013), but there are no studies on the effect of this compound on HSP70-CHIP complex formation and stability. Thus, although our data show that HSP70 inhibition did not affect CHIP migration to the nucleolus, further studies are needed to elucidate this mechanism. It may be questioned whether the accumulation of CHIP in the nucleolus implies that it is the HSP70 substrate that undergoes chaperone protection and is refolded during regeneration before being released from this compartment. Our FRAP analysis showed that approximately 30% of total CHIP was immobile in the nucleolus in the HeLa EGFP-CHIP cells. Heat shock also induces a similar formation of the immobile GFP-NPM1 protein fraction, which implies altered properties of the GC due to its association with misfolded proteins that accumulate in this phase upon heat shock (Frottin et al., 2019). Thus, it is likely that CHIP embedded in GC associates with aggregated proteins, which affects its mobility.

What is the role of CHIP in the nucleolus? We hypothesized that in collaboration with HSP70, CHIP might serve as a ubiquitin ligase or co-chaperone that regulates ubiquitination or substrate reassembly to aid in the regeneration process. To revise this, we focused on recovering a specifically modified luciferase that contained a nucleus-targeting sequence to facilitate its accumulation in the nucleolus upon heat shock. The effect of CHIP on luciferase status during heat shock and recovery but not in association with the nucleolus was previously studied *in vitro* and *in cellulo*, showing ambiguous results. CHIP can maintain denatured luciferase in a state capable of folding and ubiquitinate it *in vitro* (Rosser et al., 2007). Moreover, heat shock may enhance CHIP chaperone activity and its ability to suppress luciferase aggregation *in vitro* (Rosser et al., 2007). In heat-stressed HEK293 cells, it was demonstrated that CHIP overexpression protected luciferase activity and did not cause its increased degradation. CHIP was also able to specifically interact with thermally denatured luciferase rather than with the refolded one (Rosser et al., 2007). In fibroblasts, CHIP overexpression did not affect luciferase degradation after heat shock and during recovery but increased its HSP70-dependent reassembly and protected it from heat-induced insolubility (Kampinga et al., 2003). On the other hand, there are also conflicting data indicating that CHIP overexpression can inhibit the renaturation of denatured luciferase in Cos-7 cells and reduce HSP70 or HSP70:HSP40-mediated luciferase folding *in vitro* (Ballinger et al., 1999; Marques et al., 2006). Overall, the effect of CHIP on luciferase status is indisputable but may depend on multiple factors.

Our results suggest that CHIP abundance in nucleoli slows the rate of luciferase recovery from heat shock. Knowing that there is no increased aggregation of CHIP in nucleoli, we consider it unlikely that the presence of CHIP imposes additional stress on this organelle. Furthermore, noting that the CHIP HSP70-binding deficient K30A mutant does not significantly delay luciferase refolding despite its presence in nucleoli, we propose that CHIP controls this process via interaction with HSP70. Regulation of HSP70 by CHIP in the nucleolus may involve a reduction in the affinity of this chaperone for substrates, as shown previously (Ballinger et al., 1999; Stankiewicz et al., 2010). Also, CHIP may affect the rate of ATP hydrolysis by HSP70 (Stankiewicz et al., 2010), which may also shape the condensation state in the nucleolus and thus the environment for the recovery processes (Yewdall et al., 2021). Noteworthy, HSP70 may inhibit CHIP ubiquitination activity (Narayan et al., 2015; Das et al., 2021), resulting in a functional co-regulation of these proteins to select the optimal heat stress response in the nucleolus. We would also like to point out that we observed significantly higher CHIP levels in the nucleoli of heat-stressed HeLa EGFP-CHIP or MCF7 cells than in the luciferase-expressing HEK293T cells upon CHIP overexpression. We speculate that these particular cancer cells may more intensively utilize CHIP to manage proteostasis in the nucleus.

The role of CHIP ubiquitination activity in protein recovery in the nucleolus remains elusive and thus requires further study, perhaps by using other inactive CHIP variants and investigating their effects on ubiquitination and recovery of specific endogenous proteins. However, our FRAP analysis and data obtained from prolonged heat shock revealed altered dynamics and pro-aggregation characteristics of the catalytically inactive CHIP H260Q mutant, which we hypothesize may indirectly affect nucleolar protein regeneration. It would be interesting to investigate whether other CHIP inactive variants, also pathogenic, show greater sensitivity to heat stress, affecting nucleoli regeneration.

The mechanisms that trigger CHIP clearance from the nucleolus during regeneration are also unclear. We observed that the signals indicating the presence of luciferase and CHIP in the nucleolus decreased with similar dynamics, suggesting that some level of recovery must be achieved before CHIP is released. Recent proteomic data revealed proteins associated with NPM1 in recovering from heat shock VER-treated HEK293T cells, showing a persistent impairment of nucleolar regeneration in the presence of the inhibitor (Frottin et al., 2019). Intriguingly, CHIP was not identified in this study, although it was detected in nucleoli after heat shock when VER was not added. This may suggest that CHIP acts specifically and targets selected nucleolar proteins during the regeneration process. When we treated cells with VER throughout the heat shock and recovery, we encountered increased CHIP levels in HeLa EGFP-CHIP and MCF7 cells. This further supports our hypothesis on the functional cross-talk between CHIP and HSP70, and HSP70 inhibition-dependent CHIP response to alterations in the recovery efficiency.

We also found that cells can clear nucleolar luciferase foci even after prolonged heat shock, and this process is also affected by CHIP overexpression. Frottin et al. showed that prolonged heat shock overloads nucleolar capacity in the same cells and may be responsible for aberrant phase behavior associated with the danger of irreversible protein aggregation (Frottin et al., 2019). The process was entirely reversible in our hands, suggesting that cells adapted to stress. It is important to note that in our experimental scheme, cells were exposed to 42°C heat shock instead of 43°C. Therefore, it would be interesting to test if we balanced around the “point of no return” where we came across a temperature-dependent differential cell capability to manage stress. In conclusion, we predict that the presence of CHIP in nucleoli may provide a mechanism for selective and regulated recovery of proteins, which may be relevant for cell survival during proteotoxic stress.

## SUPPLEMENTARY FIGURE LEGENDS

Figure S1. **CHIP localizes to nucleoli specifically during heat shock**

A) A simplified scheme of genomic elements after targeted integration of the EGFP-CHIP transgene into HeLa Flp-In T-REx and 293 Flp-In T-REx cell lines (modified from (Szczesny et al., 2018)). Generated cells are resistant to hygromycin B but sensitive to zeocin selection antibiotic. The EGFP-CHIP expression is repressed by the activity of the repressor protein TetR and induced by the addition of tetracycline to the culture medium.

B) Other tested stressors do not induce CHIP migration to nucleoli. HEK EGFP-CHIP cells were exposed to various stressors: 90 min heat shock at 42°C, 50 µM sodium arsenite, 100 nM thapsigargin, 0.6 M sorbitol and 2 mg/ml puromycin for 2 h. Confocal images and their magnified views of cells after each stress are shown. The arrowhead shows CHIP in the nucleolus of the heat-stressed cell. Scale bars represent 10 µm or 5 µm (magnified views).

C) Confocal images of MCF7 cells transiently expressing EGFP-CHIP. Cells treated with heat shock show CHIP nucleolar accumulation. The lack of Hoechst 333412 staining recognizes nucleoli. A scale bar represents 5 µm.

D) Confocal images of MCF7 cells after immunostaining for CHIP (green) and NPM1 (red). Nuclei (blue) are labelled with DAPI. Images show representative cells during control conditions, after 90 min heat shock and after heat shock and recovery in the presence of the 40 µM VER inhibitor (HS + Rec + VER). Nucleoli are indicated by arrowheads and are also marked with dashed circles. A scale bar represents 5 µm.

E) Quantification of relative mean CHIP intensities: from nucleoli *vs*. nuclei in MCF7 cells from confocal images. Selected images are shown in Figure S1D. Cells were treated with heat shock and 2 h recovery without or with VER, which was applied either for complete treatment (HS + Rec + VER) or only during recovery (Rec + VER). Control cells also received 40 µM VER for 2 h. Plotted data show individual experiments. For statistical comparison a one-way *ANOVA* followed by Tukey’s multiple comparison test was used (\*\**P* < 0.01, \**P* < 0.05).

Figure S2. **Validation of HSP70 knockdown efficiency in HeLa EGFP-CHIP cells**

A) Confocal images of HeLa EGFP-CHIP cells after HSP70 knockdown *via* siRNA. To control for the effects of siRNA delivery, nontargeting siRNA was used (sineg). 72 h after siRNA transfection, control and heat-shocked cells were immunostained for HSP70 (red) and the nuclei were stained with DAPI. Scale bars represent 10 µm.

Figure S3. **Actinomycin D (Act D) alters nucleolar morphology resulting in prolonged sequestration of CHIP in nucleoli during recovery from heat shock**

A) Confocal images of HeLa EGFP-CHIP cells after immunostaining for NPM1 (left panel) and FBL (right panel). Indicated cells were pretreated with 0.05 µg/ml Act D for 2 h followed by immunostaining and confocal imaging. Arrowheads point at FBL nucleolar caps formed in the presence of Act D. Scale bars represent 10 µm.

B) Analysis of changes in nucleolar morphology upon Act D treatment. HeLa EGFP-CHIP cells were treated with 0.05 µg/ml Act D for 30 min and 2 h alone or followed by 90 min heat shock. Next, cells were fixed and immunostained for NPM1 and FBL nucleolar proteins. The mean nucleolar area, circularity (based on NPM1 signal) and percentage of nuclei with nucleolar caps (based on FBL signal) were quantified. Data are means of two (area, circularity) and three (nucleolar caps) independent experiments. Error bars represent SD. For statistical comparison a one-way *ANOVA* followed by Dunnett’s multiple comparisons test was used (\*\*\**P* < 0.001, \*\**P* < 0.01).

C) Confocal images of HeLa EGFP-CHIP cells after the 2 h-recovery from heat shock (Recovery) or pretreated with 0.05 µg/ml Act D for 2 h before the heat shock and recovery periods (Act D + Recovery). After treatment, cells were imaged live. Arrowheads indicate nucleolar CHIP in Act D-treated cells. Scale bar represents 10 µm.

Figure S4. **CHIP overexpression affects the nucleolar luciferase recovery**

A) HEK293T cells permanently expressing luciferase were transfected with vectors encoding for mCherry, mCherry-CHIP WT, mCherry-CHIP H260Q and mCherry-CHIP K30A. 24 h after transfection cells were subject to 2 h heat shock and the recovery was monitored for 6 h afterward. Cells were imaged live using confocal microscopy. Luciferase foci were counted in untransfected and mCherry-expressing cells (control groups) and cells expressing the appropriate CHIP variant. The percentage of cells with nucleolar luciferase foci was determined for each condition. Data are means of three independent experiments. Error bars represent SD. For statistical comparison a two-way *ANOVA* with post hoc Tukey’s test was used (\*\*\*\**P* < 0.0001, \*\*\**P* < 0.001, \*\**P* < 0.01, \**P* < 0.05).

B) HEK293T cells permanently expressing luciferase were transfected with vectors encoding for mCherry, mCherry-CHIP WT, mCherry-CHIP H260Q, and mCherry-CHIP K30A. Transfection was performed with 0.2 µg plasmids per well of the 8-well chamber. 24 h after transfection cells were subject to 2 h-heat shock followed by 2 h-recovery. Images of live cells at the indicated time were taken using confocal microscopy. Luciferase foci were counted in untransfected and mCherry-expressing cells (control groups) and cells expressing the appropriate CHIP variant. The percentage of cells with nucleolar luciferase foci was determined for each condition. Data normalization was performed as described in Fig. 6B. Data are means of three independent experiments. Error bars represent SD. For statistical comparison, a two-way *ANOVA* with post hoc Tukey’s test was used \*\*\**P* < 0.001, \*\**P* < 0.01, \**P* < 0.05).

Figure S5. **During prolonged heat shock, luciferase recovery in nucleoli is affected by CHIP WT and CHIP H260Q overexpression**

A) HEK293T cells permanently expressing luciferase were transfected with vectors encoding for mCherry, mCherry-CHIP WT, mCherry-CHIP H260Q and mCherry-CHIP K30A and treated with 6 h heat shock. Every 2 h cells were imaged live using confocal microscopy. Images were analyzed for mean intensities of GFP-tagged luciferase in whole nucleoli relative to nuclei. Data are means of three independent experiments. Error bars represent SD. For statistical comparison a two-way *ANOVA* with post hoc Tukey’s test was used (\*\*\*\**P* < 0.0001, \*\*\**P* < 0.001, *ns P* > 0.05).

B) HEK293T cells permanently expressing luciferase were transfected with vectors encoding for mCherry, mCherry-CHIP WT, mCherry-CHIP H260Q and mCherry-CHIP K30A and treated with 6 h heat shock. During treatments cells were imaged by confocal microscopy. Luciferase foci were counted in untransfected and mCherry-expressing cells (control groups) and cells expressing the appropriate CHIP variant. The percentages of total cell counts were quantified for each condition. Data normalization was performed as described in Fig. 6B. Data are means of three independent experiments. Error bars represent SD. For statistical comparison, a two-way *ANOVA* with post hoc Tukey’s test was used (\*\*\**P* < 0.001, *ns P* > 0.05).

C) CHIP WT and H260Q show sustained sequestration into nucleoli during prolonged heat stress. HEK293T cells permanently expressing luciferase were transfected with vectors encoding for mCherry-CHIP WT, mCherry-CHIP H260Q and mCherry-CHIP K30A and treated with 6 h heat shock. Every 2 h intervals, confocal images of live cells were taken to measure CHIP intensities in nucleoli and nuclei, and the relative intensities were quantified. Data are means of three independent experiments. Error bars represent SD. For statistical comparison a two-way *ANOVA* with post hoc Tukey’s test was used (\*\*\*\**P* < 0.0001, \*\*\**P* < 0.001, \*\**P* < 0.01, \**P* < 0.05, *ns P* > 0.05).

## MATERIALS AND METHODS

### METHODS

#### Cell culture

HeLa Flp-In T-REx, HEK293 Flp-In T-REx (a kind gift from Dr. R. Szczesny), MCF7 (a kind gift from Prof. A. Zylicz), HeLa EGFP-CHIP, HEK EGFP-CHIP, and HEK293T NLS LG cells (stably expressing luciferase; a kind gift from Dr. M.S. Hipp) were cultured in Dulbecco’s Modified Eagle’s Medium (D6429, Sigma) supplemented with 10% heat-inactivated fetal bovine serum (Sigma) and 1% antibiotic - antimycotic (Gibco) at 37°C with 5% CO_2_ in a humidified incubator. To maintain stable cell lines, HeLa EGFP-CHIP and HEK EGFP-CHIP cells were supplemented with blasticidin (10 µg/ml) (ant-bl-1, Invivogen) and hygromycin B (50 µg/ml) (10687010, Thermo Fisher Scientific), while 293T NLS LG cells were supplemented with G418 (100 µg/ml) (10131035, Gibco). For the experiment, the EGFP-CHIP expression in HeLa EGFP-CHIP and HEK EGFP-CHIP cells was induced by adding tetracycline to the medium (25 ng/ml) upon plating. Where indicated, to induce heat shock, cells were transferred to another humidified incubator set at 42°C. For the recovery, cells were transferred back to 37°C. For passaging and experiments, cells were dissociated from the plate with trypsin (Trypsin-EDTA 0.25%, Sigma). Cells were tested for mycoplasma using a PCR-based assay.

#### Poly-L-lysine coating

HEK293T cells were grown on cover glasses or 8 well-chambered slides (Ibidi) coated with Poly-L-lysine solution (P4707, Sigma). The coating was performed for 1 h at 37°C, followed by two washes in sterile PBS and drying under a laminar flow hood.

#### Tetracycline preparation

Tetracycline was prepared according to an established protocol (Szczesny et al., 2018). Briefly, tetracycline was added to 96% ethanol at the 5 mg/ml concentration. The solution was rotated for 30 min at room temperature and incubated overnight at -20°C. The next day the rotation was repeated for 30 min. Afterward, it was filtered through the 0.22 μm syringe filter and diluted with ethanol to the final concentration of 100 μg/ml. The solution was stored at -20°C.

#### Plasmid construction

Vectors: pKK-EGFP-TEV and pKK-mCherry-TEV were a kind gift from Dr. R. Szczesny.

The sequence and ligation independent cloning (SLIC) method was used to construct mCherry-CHIP, mCherry-CHIP H260Q, mCherry-CHIP K30A, EGFP-CHIP, EGFP-CHIP H260Q plasmids. The parental vectors (pKK-EGFP-TEV and pKK-mCherry-TEV) were linearized with BshTI i NheI enzymes. For SLIC cloning, linearized vectors were mixed with PCR-amplified human CHIP sequence and treated with T4 DNA polymerase, followed by bacterial transformation. The following primers were used for CHIP sequence amplification:

**Hchip forward**:

GGATCCgaaaacctgtacttccaaggaACCGGTATGAAGGGCAAGGAGGAGAAG

**hCHIP reverse**:

GATATCaccctgaaaatacaaattctcGCTAGCTCAGTAGTCCTCCACCCAGC

To insert H260Q mutation into the CHIP sequence, two PCR reactions were carried out using the following primers:

The 1^st^ amplicon:

**hCHIP forward**:

GGATCCgaaaacctgtacttccaaggaACCGGTATGAAGGGCAAGGAGGAGAAG

**r1**: CACGCTGCAGcTGCTCCTCGATGTCC

The 2^nd^ amplicon:

**f1**: GGACATCGAGGAGCAgCTGCAGCGTG

**hCHIP reverse**:

GATATCaccctgaaaatacaaattctcGCTAGCTCAGTAGTCCTCCACCCAGC

Afterward, splice-PCR was used to assemble both fragments.

Sequence validation was performed using restriction enzymes (BamHI and EcoRV) and sequencing with the following primers:

**FRTTO_For** tgacctccatagaagacacc

**FRTTO_Rev** aactagaaggcacagtcgag

**EGFP_F** catggtcctgctggagttcg

**CHIP_For** atgaagggcaaggaggagaag

To generate EGFP-CHIP K30A plasmid, Q5 Site-Directed Mutagenesis Kit (E0554S, New England Biolabs) was used with the following primers:

**F-hCHIP K30A**: GCAGGAGCTCgcGGAGCAGGGCAATC

**R-hCHIP K30A**: GCGCTCGGGCTCTTCTCG

#### Stable cell line generation

HeLa Flp-In T-REx and HEK293 Flp-In T-REx cells were grown on 6-well plates. For stable cell line generation, the cells were co-transfected with 1 μg pOG44 (a kind gift from Dr. R. Szczesny) and 0.8 μg EGFP-CHIP plasmid using Mirus reagents: 2 ul TransIT-293 (MIR 2700, Mirus) for transfection of HEK293 Flp-In T-REx cells and 2 ul Trans-IT-HeLa and 1.3 ul Monster (MIR 2900, Mirus) for HeLa Flp-In T-REx cells. The day following transfection cells were treated with selection antibiotics: 10 μg/ml blasticidin (Invivogen) and 50 μg/ml hygromycin B (Thermo Fisher Scientific). The treatment continued for a month.

#### Cell transfection

Transient transfections were performed using Lipofectamine 2000 (Invitrogen) according to the manufacturer’s guidelines. Cells seeded on 8 well-chambered slides (Ibidi) were transfected with 0.5 μg plasmid per well. Cells seeded for VER-155008 treatment were transfected with 0.2 μg plasmid. Transfections were carried out a day before imaging.

#### HSP70 knockdown

HeLa EGFP-CHIP cells were seeded on 35 mm imaging dishes (Ibidi). The following day cells were co-transfected with 75 pmols HSP70 siRNAs (IDs 145248 and 6965, Thermo Fisher Scientific) using 9 μl Lipofectamine RNAiMAX reagent (Invitrogen) in Opti-MEM Reduced Serum Medium. The medium was exchanged after 48 h post-transfection. Silencing lasted 72 h.

#### Actinomycin D and VER-155008 treatments

Cells were treated with 0.05 µg/ml Actinomycin D (Act D) (1229, Tocris) dissolved in DMSO in a complete medium for 0.5 or 2 h at 37°C. Then the cells were exposed to 90 min heat shock at 42°C and 120 min of recovery at 37°C. The cells were fixed for immunofluorescence, or live cells were imaged using confocal microscopy.

VER-155008 (SML0271, Sigma) dissolved in DMSO was added to the complete medium at the final concentration of 40 μM. HEK293T and MCF7 cells were treated with VER-155008 before the heat shock, while HeLa EGFP-CHIP cells were pretreated with VER-155008 for 2 h before the heat shock. All cell lines were treated with VER-155008 immediately after the heat shock for recovery. Cells were fixed for immunofluorescence, or live cells were imaged using confocal microscopy.

#### Immunofluorescence

Cells were fixed with 4% (para)formaldehyde (28906, Pierce) in PBS for 10 min at room temperature before washing 3 times with PBS for 5 min. The cells were permeabilized for 10 min with Triton X-100 0.1% (v/v) in PBS at room temperature. Samples were incubated in a blocking buffer (either 2% bovine serum albumin BSA, 1.5% goat serum, 0.1% Triton X-100 in PBS, or 1% BSA in PBS) for 10 min at room temperature. Primary antibodies were applied in a blocking buffer and incubated overnight at 4°C. Appropriate fluorescent secondary antibodies at a dilution of 1:500 were applied after PBS washes for 60 min at room temperature in PBS. Samples were mounted using the Vectashield antifade mounting medium with DAPI (Vector Laboratories).

Primary antibodies:

▪ CHIP (1:250) rabbit [EPR4447] (ab134064) Abcam
▪ NPM1 (1:250) mouse (32-5200) Invitrogen
▪ FBL (1:400) rabbit (2639S) Cell Signaling Technology
▪ HSP70 (1:250) mouse (SMC-100) StressMarq Biosciences

Secondary antibodies:

Goat Alexa 647 anti-mouse (A21235, Invitrogen), goat Alexa 568 anti-mouse (A11031, Invitrogen), goat Alexa 647 anti-rabbit (A21245, Invitrogen).

#### Image acquisition

Confocal microscopy was performed on the ZEISS LSM800 confocal laser scanning microscope (Carl Zeiss Microscopy) using 63x/1.4 NA or 40x/1.3 NA oil immersion objectives. Images show single optical sections. Within each experiment, images were acquired using identical acquisition settings. For colocalization studies, imaging was carried out with Airyscan.

#### Image analysis

ImageJ software (https://imagej.nih.gov/ij/index.html) (Schneider et al., 2012) was used for image analysis in most experiments except for MCF7 immunostaining. In HeLa EGFP-CHIP cells, nucleoli were manually selected, and the CHIP (EGFP) intensity (mean gray value) was calculated. For colocalization studies, the JaCoP plugin was used (Bolte and Cordelières, 2006). The image background was corrected using the rolling ball algorithm (rolling ball=150). Thresholds of the green and red channels were selected manually and maintained in every image.

#### Analysis of the fluorescence intensity ratio - nucleolus: nucleus in HEK293T cells

Nucleoli were manually located using the Hoechst 33342 channel. Relative luciferase or CHIP concentrations in nucleoli/nuclei were calculated based on GFP or mCherry intensities, respectively, in each compartment in 50 cells per condition across three biological repeats.

#### Quantification of nucleolar luciferase foci in HEK293T cells

Luciferase foci were counted in 47-142 cells (66-301 cells for the experiments with VER-155008) per time point and condition across 3 biological repeats.

#### Analysis of CHIP ratio in MCF7 cells

Image analysis was conducted with a customized CellProfiler 4.2.1. (Carpenter et al., 2006; Stirling et al., 2021) pipeline. In short, nuclei and nucleoli objects were segmented from DAPI and NPM1 channels, respectively, using the three-class Otsu thresholding method, excluding objects touching the image’s border. Next, the nuclei objects were masked by nucleoli objects, thus creating the third class of objects – masked nuclei, consisting solely of nucleoplasm without nucleoli. The relationship of each nucleolus object to its parent nucleus object was assigned using the RelateObjects module. Finally, CHIP intensity was calculated from the CHIP channel for both masked nuclei and nucleoli objects and exported together with the relationship information to CSV files. For each repetition of the experiment, the ratio of each child nucleolus/parent masked nucleus mean intensities was calculated. Nucleoli without assigned parent nucleus (parent ID 0) were discarded from the analysis.

#### FRAP

Cells used in FRAP studies were cultured on 35-mm imaging dishes (Ibidi). FRAP experiments were performed on ZEISS LSM800 confocal laser scanning microscope equipped with the 40x/1.3 NA oil immersion objective. A circular region of interest of the constant size was selected within nucleoli, and bleaching was carried out with 100% laser power of the 488 nm laser line. Fluorescence intensity was recorded for up to 3 min at a frame interval of 0.5 s. FRAP movies were analyzed using FRAPAnalyser (https://github.com/ssgpers/FRAPAnalyser). Fluorescence intensity was corrected for background fluorescence and photobleaching. Recovery curves were fitted with a single exponential recovery.

#### Nucleoli isolation

Cells grown on 100 mm tissue culture dishes were harvested on ice. The isolation of nucleoli was performed according to the previously described protocol with some modifications (Andersen et al., 2002). The old medium was discarded, and the cells were washed 1× with 4 ml of ice-cold PBS. Cells were harvested on ice in cold PBS. The cells were washed 1× with ice-cold PBS at 220 × g at 4°C. After the PBS wash, the cells were resuspended in 5 ml of Buffer A (10 mM HEPES pH 7.9, 10 mM KCl, mM MgCl_2_, 0.5 mM DTT, 1x Complete protease inhibitor cocktail (Roche)) and incubated on ice for 5 min. The cells were homogenized with a pre-cooled 1 ml Dounce homogenizer (Wheaton) on ice 10× using a tight pestle. The homogenized cells were centrifuged at 220 × g for 5 min at 4°C. The supernatant was collected as the cytoplasmic fraction. The pellet was resuspended with 3 ml S1 solution (0.25 M sucrose, 10 mM MgCl_2_, 1x Complete protease inhibitor cocktail (Roche)) by pipetting up and down. The resuspended pellet was layered carefully over 3 ml of S2 solution (0.35 M sucrose, 0.5 mM MgCl_2_, 1x Complete protease inhibitor cocktail (Roche)) and centrifuged at 1430 × g for 5 min at 4°C. The pellet was resuspended with 3 ml of S2 solution by pipetting up and down. The nuclear suspension was sonicated on the ice at 50% power 11×, each time for 10 s and 10 s of rest on ice (Sonica VCX130 with a ¼ inch tip). The sonicated sample was layered over 3 ml of S3 solution (0.88 M sucrose, 0.5 mM MgCl_2_, 1x Complete protease inhibitor cocktail (Roche)) in a new Falcon tube and centrifuged at 3000 × g for 10 min at 4°C. The supernatant was collected as the nucleoplasm fraction. The pellet was resuspended with 500 µl of S2 solution and centrifuged at 1430 × g for 5 min at 4°C. The nucleoli were resuspended in 500 µL of S2 solution and stored at –80°C as nucleoli fractions.

#### Estimation of the protein concentration and Western blotting

The protein concentration was estimated using the BCA protein assay kit (23225, Thermo Scientific). Protein samples in SDS-loading dye (reducing) were run in 10% acrylamide gels in a running buffer (25 mM Tris, 190 mM glycine, 0.1% SDS) at 90 V (stacking gel) and 150 V (separating gel). The wet transfer was done at a constant 400 mA for 1 h at 4°C in a transfer buffer (25 mM Tris, 190 mM glycine, 20% methanol, pH 8.3). Blots were blocked with 5% skimmed milk in TBST (50 mM Tris, 150 mM NaCl, 0.1% Tween 20, pH 7.6) for 1 h at room temperature and incubated overnight with primary antibodies prepared in 5% skimmed milk in TBST at 4°C. The blots were then washed three times with TBST for 10 min each wash. Finally, the blots were incubated with horseradish peroxidase-linked secondary antibodies (1:10000) prepared in 5% skimmed milk in TBST for 1 h at room temperature. Imaging was performed using a ChemiDoc™ Imaging System (Bio-Rad).

Antibodies:

▪ CHIP (1: 1000) rabbit [EPR4447] (ab134064) Abcam
▪ FBL (1: 500) rabbit (2639S) Cell Signaling Technology
▪ Alpha-tubulin (1: 1000) mouse (32-2500) Invitrogen
▪ Lamin B1 (1: 1000) mouse (33-2000) Invitrogen
▪ HSP70 (1: 500) mouse (SMC-100) StressMarq Biosciences

#### *In vitro* ubiquitination assay

The reactions were run at 37°C for 90 min using 60 µM Ubiquitin (Boston Biochem) in the presence of 100 nM E1 (UBE1, Boston Biochem), 0.6 µM E2 (Boston Biochem), E3 ligase reaction buffer (Boston Biochem), and ATP in 25 μl reaction mixture. 2 μl cell lysates served as a source of the E3 ligase CHIP (cells were lysed in Cell lysis buffer (9803, Cell Signaling Technology). After reactions, protein samples were mixed with the SDS-loading dye and boiled for Western blot analysis.

#### Statistical analysis

Data were plotted and analyzed with the GraphPad Prism 9 software. P-values were calculated using a two-way or one way *ANOVA* followed by multiple comparisons tests. Two-tailed unpaired t-test or Mann-Whitney test were used to compare differences between two independent groups. In the figures, * = p< 0.05, ** = p< 0.01, *** = p< 0.001, **** = p< 0.0001.

## ACKNOWLEDGEMENTS

We thank the Genome Engineering Unit of the International Institute of Molecular and Cell Biology in Warsaw for generating DNA constructs. We thank the Microscopy and Cytometry Facility of the International Institute of Molecular and Cell Biology in Warsaw for assistance with the confocal microscopy. We thank Aleksandra Szybińska for optimizing the HSP70 knockdown in HeLa EGFP-CHIP cells. We thank members of the Pokrzywa laboratory for discussions and comments on the manuscript.

## FUNDING

L.B., K.K. and W.P. were supported by the Foundation for Polish Science, co-financed by the European Union under the European Regional Development Fund (grant POIR.04.04.00-00-5EAB/18-00) M.P. was supported by the Norwegian Financial Mechanism 2014-2021 and operated by the Polish National Science Center under the project contract no UMO-2019/34/H/NZ3/00691.

## CONFLICT OF INTEREST

The authors declare that the research was conducted without any commercial or financial relationships that could be construed as a potential conflict of interest.

## AUTHOR CONTRIBUTIONS

Contributions of individual authors based on the [CRediT](https://casrai.org/credit/) (Contributor Roles Taxonomy).

**Malgorzata Piechota:** Conceptualization; Data curation; Formal analysis; Methodology; Investigation; Validation; Visualization; Supervision; Validation; Writing-original draft; Writing-review & editing. **Lilla Biriczova:** Formal analysis; Investigation; Methodology; Visualization; Writing-review & editing. **Konrad Kowalski**: Formal analysis; Investigation; Methodology; Visualization; Writing-review & editing. **Natalia A. Szulc**: Formal analysis; Writing-review & editing. **Wojciech Pokrzywa:** Conceptualization; Data curation; Formal analysis; Funding acquisition; Project administration; Resources; Supervision; Validation; Writing-original draft; Writing-review & editing.

## Notes

### Competing Interest Statement

The authors have declared no competing interest.

## REFERENCES

Alberti, S., and Carra, S. (2019). Nucleolus: A Liquid Droplet Compartment for Misbehaving Proteins. Curr Biol 29(19), R930–r932. doi: 10.1016/j.cub.2019.08.013.

Andersen, J.S., Lyon, C.E., Fox, A.H., Leung, A.K., Lam, Y.W., Steen, H., et al. (2002). Directed proteomic analysis of the human nucleolus. Curr Biol 12(1), 1–11. doi: 10.1016/s0960-9822(01)00650-9.

Audas, T.E., Jacob, M.D., and Lee, S. (2012). Immobilization of proteins in the nucleolus by ribosomal intergenic spacer noncoding RNA. Mol Cell 45(2), 147–157. doi: 10.1016/j.molcel.2011.12.012.

Azkanaz, M., Rodríguez López, A., de Boer, B., Huiting, W., Angrand, P.O., Vellenga, E., et al. (2019). Protein quality control in the nucleolus safeguards recovery of epigenetic regulators after heat shock. Elife 8. doi: 10.7554/eLife.45205.

Ballinger, C.A., Connell, P., Wu, Y., Hu, Z., Thompson, L.J., Yin, L.Y., et al. (1999). Identification of CHIP, a novel tetratricopeptide repeat-containing protein that interacts with heat shock proteins and negatively regulates chaperone functions. Mol Cell Biol 19(6), 4535–4545. doi: 10.1128/mcb.19.6.4535.

Biggiogera, M., Bürki, K., Kaufmann, S.H., Shaper, J.H., Gas, N., Amalric, F., et al. (1990). Nucleolar distribution of proteins B23 and nucleolin in mouse preimplantation embryos as visualized by immunoelectron microscopy. Development 110(4), 1263–1270. doi: 10.1242/dev.110.4.1263.

Bolte, S., and Cordelières, F.P. (2006). A guided tour into subcellular colocalization analysis in light microscopy. J Microsc 224(Pt 3), 213–232. doi: 10.1111/j.1365-2818.2006.01706.x.

Buetow, L., and Huang, D.T. (2016). Structural insights into the catalysis and regulation of E3 ubiquitin ligases. Nat Rev Mol Cell Biol 17(10), 626–642. doi: 10.1038/nrm.2016.91.

Carpenter, A.E., Jones, T.R., Lamprecht, M.R., Clarke, C., Kang, I.H., Friman, O., et al. (2006). CellProfiler: image analysis software for identifying and quantifying cell phenotypes. Genome Biol 7(10), R100. doi: 10.1186/gb-2006-7-10-r100.

Cerqueira, A.V., and Lemos, B. (2019). Ribosomal DNA and the Nucleolus as Keystones of Nuclear Architecture, Organization, and Function. Trends Genet 35(10), 710–723. doi: 10.1016/j.tig.2019.07.011.

Dai, Q., Zhang, C., Wu, Y., McDonough, H., Whaley, R.A., Godfrey, V., et al. (2003). CHIP activates HSF1 and confers protection against apoptosis and cellular stress. Embo j 22(20), 5446–5458. doi: 10.1093/emboj/cdg529.

Das, A., Thapa, P., Santiago, U., Shanmugam, N., Banasiak, K., Dabrowska, K., et al. (2021). Heterotypic Assembly Mechanism Regulates CHIP E3 Ligase Activity. bioRxiv, 2021.2008.2020.457118. doi: 10.1101/2021.08.20.457118.

Demand, J., Alberti, S., Patterson, C., and Höhfeld, J. (2001). Cooperation of a ubiquitin domain protein and an E3 ubiquitin ligase during chaperone/proteasome coupling. Curr Biol 11(20), 1569–1577. doi: 10.1016/s0960-9822(01)00487-0.

Feric, M., Vaidya, N., Harmon, T.S., Mitrea, D.M., Zhu, L., Richardson, T.M., et al. (2016). Coexisting Liquid Phases Underlie Nucleolar Subcompartments. Cell 165(7), 1686–1697. doi: 10.1016/j.cell.2016.04.047.

Frottin, F., Schueder, F., Tiwary, S., Gupta, R., Körner, R., Schlichthaerle, T., et al. (2019). The nucleolus functions as a phase-separated protein quality control compartment. Science 365(6451), 342–347. doi: 10.1126/science.aaw9157.

Hatakeyama, S., Yada, M., Matsumoto, M., Ishida, N., and Nakayama, K.I. (2001). U box proteins as a new family of ubiquitin-protein ligases. J Biol Chem 276(35), 33111–33120. doi: 10.1074/jbc.M102755200.

Huang, S. (2002). Building an efficient factory: where is pre-rRNA synthesized in the nucleolus? J Cell Biol 157(5), 739–741. doi: 10.1083/jcb.200204159.

Jiang, J., Ballinger, C.A., Wu, Y., Dai, Q., Cyr, D.M., Höhfeld, J., et al. (2001). CHIP is a U-box-dependent E3 ubiquitin ligase: identification of Hsc70 as a target for ubiquitylation. J Biol Chem 276(46), 42938–42944. doi: 10.1074/jbc.M101968200.

Jordan, E.G. (1984). Nucleolar nomenclature. J Cell Sci 67, 217–220. doi: 10.1242/jcs.67.1.217.

Joshi, V., Amanullah, A., Upadhyay, A., Mishra, R., Kumar, A., and Mishra, A. (2016). A Decade of Boon or Burden: What Has the CHIP Ever Done for Cellular Protein Quality Control Mechanism Implicated in Neurodegeneration and Aging? Front Mol Neurosci 9, 93. doi: 10.3389/fnmol.2016.00093.

Kampinga, H.H., Kanon, B., Salomons, F.A., Kabakov, A.E., and Patterson, C. (2003). Overexpression of the cochaperone CHIP enhances Hsp70-dependent folding activity in mammalian cells. Mol Cell Biol 23(14), 4948–4958. doi: 10.1128/MCB.23.14.4948-4958.2003.

Komander, D. (2009). The emerging complexity of protein ubiquitination. Biochem Soc Trans 37(Pt 5), 937–953. doi: 10.1042/bst0370937.

Kozakai, Y., Kamada, R., Furuta, J., Kiyota, Y., Chuman, Y., and Sakaguchi, K. (2016). PPM1D controls nucleolar formation by up-regulating phosphorylation of nucleophosmin. Sci Rep 6, 33272. doi: 10.1038/srep33272.

Krüger, T., Zentgraf, H., and Scheer, U. (2007). Intranucleolar sites of ribosome biogenesis defined by the localization of early binding ribosomal proteins. J Cell Biol 177(4), 573–578. doi: 10.1083/jcb.200612048.

Lafontaine, D.L.J., Riback, J.A., Bascetin, R., and Brangwynne, C.P. (2021). The nucleolus as a multiphase liquid condensate. Nat Rev Mol Cell Biol 22(3), 165–182. doi: 10.1038/s41580-020-0272-6.

Latonen, L., Moore, H.M., Bai, B., Jäämaa, S., and Laiho, M. (2011). Proteasome inhibitors induce nucleolar aggregation of proteasome target proteins and polyadenylated RNA by altering ubiquitin availability. Oncogene 30(7), 790–805. doi: 10.1038/onc.2010.469.

Lindström, M.S., Jurada, D., Bursac, S., Orsolic, I., Bartek, J., and Volarevic, S. (2018). Nucleolus as an emerging hub in maintenance of genome stability and cancer pathogenesis. Oncogene 37(18), 2351–2366. doi: 10.1038/s41388-017-0121-z.

Marques, C., Guo, W., Pereira, P., Taylor, A., Patterson, C., Evans, P.C., et al. (2006). The triage of damaged proteins: degradation by the ubiquitin-proteasome pathway or repair by molecular chaperones. Faseb j 20(6), 741–743. doi: 10.1096/fj.05-5080fje.

Meacham, G.C., Patterson, C., Zhang, W., Younger, J.M., and Cyr, D.M. (2001). The Hsc70 co-chaperone CHIP targets immature CFTR for proteasomal degradation. Nat Cell Biol 3(1), 100–105. doi: 10.1038/35050509.

Mediani, L., Guillén-Boixet, J., Vinet, J., Franzmann, T.M., Bigi, I., Mateju, D., et al. (2019). Defective ribosomal products challenge nuclear function by impairing nuclear condensate dynamics and immobilizing ubiquitin. Embo j 38(15), e101341. doi: 10.15252/embj.2018101341.

Mekhail, K., Khacho, M., Carrigan, A., Hache, R.R., Gunaratnam, L., and Lee, S. (2005). Regulation of ubiquitin ligase dynamics by the nucleolus. J Cell Biol 170(5), 733–744. doi: 10.1083/jcb.200506030.

Mitrea, D.M., Cika, J.A., Stanley, C.B., Nourse, A., Onuchic, P.L., Banerjee, P.R., et al. (2018). Self-interaction of NPM1 modulates multiple mechanisms of liquid-liquid phase separation. Nat Commun 9(1), 842. doi: 10.1038/s41467-018-03255-3.

Murata, S., Minami, Y., Minami, M., Chiba, T., and Tanaka, K. (2001). CHIP is a chaperone-dependent E3 ligase that ubiquitylates unfolded protein. EMBO Rep 2(12), 1133–1138. doi: 10.1093/embo-reports/kve246.

Narayan, V., Landré, V., Ning, J., Hernychova, L., Muller, P., Verma, C., et al. (2015). Protein-Protein Interactions Modulate the Docking-Dependent E3-Ubiquitin Ligase Activity of Carboxy-Terminus of Hsc70-Interacting Protein (CHIP). Mol Cell Proteomics 14(11), 2973–2987. doi: 10.1074/mcp.M115.051169.

Nollen, E.A., Salomons, F.A., Brunsting, J.F., van der Want, J.J., Sibon, O.C., and Kampinga, H.H. (2001). Dynamic changes in the localization of thermally unfolded nuclear proteins associated with chaperone-dependent protection. Proc Natl Acad Sci U S A 98(21), 12038–12043. doi: 10.1073/pnas.201112398.

Pelham, H., Lewis, M., and Lindquist, S. (1984). Expression of a Drosophila heat shock protein in mammalian cells: transient association with nucleoli after heat shock. Philos Trans R Soc Lond B Biol Sci 307(1132), 301–307. doi: 10.1098/rstb.1984.0131.

Pelham, H.R. (1984). Hsp70 accelerates the recovery of nucleolar morphology after heat shock. Embo j 3(13), 3095–3100. doi: 10.1002/j.1460-2075.1984.tb02264.x.

Petrucelli, L., Dickson, D., Kehoe, K., Taylor, J., Snyder, H., Grover, A., et al. (2004). CHIP and Hsp70 regulate tau ubiquitination, degradation and aggregation. Hum Mol Genet 13(7), 703–714. doi: 10.1093/hmg/ddh083.

Qian, S.B., McDonough, H., Boellmann, F., Cyr, D.M., and Patterson, C. (2006). CHIP-mediated stress recovery by sequential ubiquitination of substrates and Hsp70. Nature 440(7083), 551–555. doi: 10.1038/nature04600.

Reynolds, R.C., Montgomery, P.O., and Hughes, B. (1964). NUCLEOLAR “CAPS” PRODUCED BY ACTINOMYCIN D. Cancer Res 24, 1269–1277.

Riback, J.A., Zhu, L., Ferrolino, M.C., Tolbert, M., Mitrea, D.M., Sanders, D.W., et al. (2020). Composition-dependent thermodynamics of intracellular phase separation. Nature 581(7807), 209–214. doi: 10.1038/s41586-020-2256-2.

Rosser, M.F., Washburn, E., Muchowski, P.J., Patterson, C., and Cyr, D.M. (2007). Chaperone functions of the E3 ubiquitin ligase CHIP. J Biol Chem 282(31), 22267–22277. doi: 10.1074/jbc.M700513200.

Scheer, U., and Hock, R. (1999). Structure and function of the nucleolus. Curr Opin Cell Biol 11(3), 385–390. doi: 10.1016/s0955-0674(99)80054-4.

Schlecht, R., Scholz, S.R., Dahmen, H., Wegener, A., Sirrenberg, C., Musil, D., et al. (2013). Functional analysis of Hsp70 inhibitors. PLoS One 8(11), e78443. doi: 10.1371/journal.pone.0078443.

Schneider, C.A., Rasband, W.S., and Eliceiri, K.W. (2012). NIH Image to ImageJ: 25 years of image analysis. Nat Methods 9(7), 671–675. doi: 10.1038/nmeth.2089.

Shav-Tal, Y., Blechman, J., Darzacq, X., Montagna, C., Dye, B.T., Patton, J.G., et al. (2005). Dynamic sorting of nuclear components into distinct nucleolar caps during transcriptional inhibition. Mol Biol Cell 16(5), 2395–2413. doi: 10.1091/mbc.e04-11-0992.

Shimura, H., Schwartz, D., Gygi, S.P., and Kosik, K.S. (2004). CHIP-Hsc70 complex ubiquitinates phosphorylated tau and enhances cell survival. J Biol Chem 279(6), 4869–4876. doi: 10.1074/jbc.M305838200.

Smirnov, E., Borkovec, J., Kováčik, L., Svidenská, S., Schröfel, A., Skalníková, M., et al. (2014). Separation of replication and transcription domains in nucleoli. J Struct Biol 188(3), 259–266. doi: 10.1016/j.jsb.2014.10.001.

Stankiewicz, M., Nikolay, R., Rybin, V., and Mayer, M.P. (2010). CHIP participates in protein triage decisions by preferentially ubiquitinating Hsp70-bound substrates. Febs j 277(16), 3353–3367. doi: 10.1111/j.1742-4658.2010.07737.x.

Stirling, D.R., Swain-Bowden, M.J., Lucas, A.M., Carpenter, A.E., Cimini, B.A., and Goodman, A. (2021). CellProfiler 4: improvements in speed, utility and usability. BMC Bioinformatics 22(1), 433. doi: 10.1186/s12859-021-04344-9.

Szaflarski, W., Leśniczak-Staszak, M., Sowiński, M., Ojha, S., Aulas, A., Dave, D., et al. (2022). Early rRNA processing is a stress-dependent regulatory event whose inhibition maintains nucleolar integrity. Nucleic Acids Res 50(2), 1033–1051. doi: 10.1093/nar/gkab1231.

Szczesny, R.J., Kowalska, K., Klosowska-Kosicka, K., Chlebowski, A., Owczarek, E.P., Warkocki, Z., et al. (2018). Versatile approach for functional analysis of human proteins and efficient stable cell line generation using FLP-mediated recombination system. PLoS One 13(3), e0194887. doi: 10.1371/journal.pone.0194887.

Tateishi, Y., Kawabe, Y., Chiba, T., Murata, S., Ichikawa, K., Murayama, A., et al. (2004). Ligand-dependent switching of ubiquitin-proteasome pathways for estrogen receptor. Embo j 23(24), 4813–4823. doi: 10.1038/sj.emboj.7600472.

Wang, M., Bokros, M., Theodoridis, P.R., and Lee, S. (2019). Nucleolar Sequestration: Remodeling Nucleoli Into Amyloid Bodies. Front Genet 10, 1179. doi: 10.3389/fgene.2019.01179.

Welch, W.J., and Feramisco, J.R. (1984). Nuclear and nucleolar localization of the 72,000-dalton heat shock protein in heat-shocked mammalian cells. J Biol Chem 259(7), 4501–4513.

Welch, W.J., and Suhan, J.P. (1986). Cellular and biochemical events in mammalian cells during and after recovery from physiological stress. J Cell Biol 103(5), 2035–2052. doi: 10.1083/jcb.103.5.2035.

Wu, X., and Hammer, J.A. (2021). ZEISS Airyscan: Optimizing Usage for Fast, Gentle, Super-Resolution Imaging. Methods Mol Biol 2304, 111–130. doi: 10.1007/978-1-0716-1402-0_5.

Yang, K., Wang, M., Zhao, Y., Sun, X., Yang, Y., Li, X., et al. (2016). A redox mechanism underlying nucleolar stress sensing by nucleophosmin. Nat Commun 7, 13599. doi: 10.1038/ncomms13599.

Yewdall, N.A., André, A.A.M., van Haren, M.H.I., Nelissen, F.H.T., Jonker, A., and Spruijt, E. (2021). ATP:Mg2+ shapes condensate properties of rRNA-NPM1 in vitro nucleolus model and its partitioning of ribosomes. bioRxiv, 2021.2012.2022.473778. doi: 10.1101/2021.12.22.473778.

Younger, J.M., Ren, H.Y., Chen, L., Fan, C.Y., Fields, A., Patterson, C., et al. (2004). A foldable CFTR{Delta}F508 biogenic intermediate accumulates upon inhibition of the Hsc70-CHIP E3 ubiquitin ligase. J Cell Biol 167(6), 1075–1085. doi: 10.1083/jcb.200410065.

