## Supplementary figures and images for "CHIP ubiquitin ligase is involved in the nucleolar stress management"

### Supplemental Figure 1

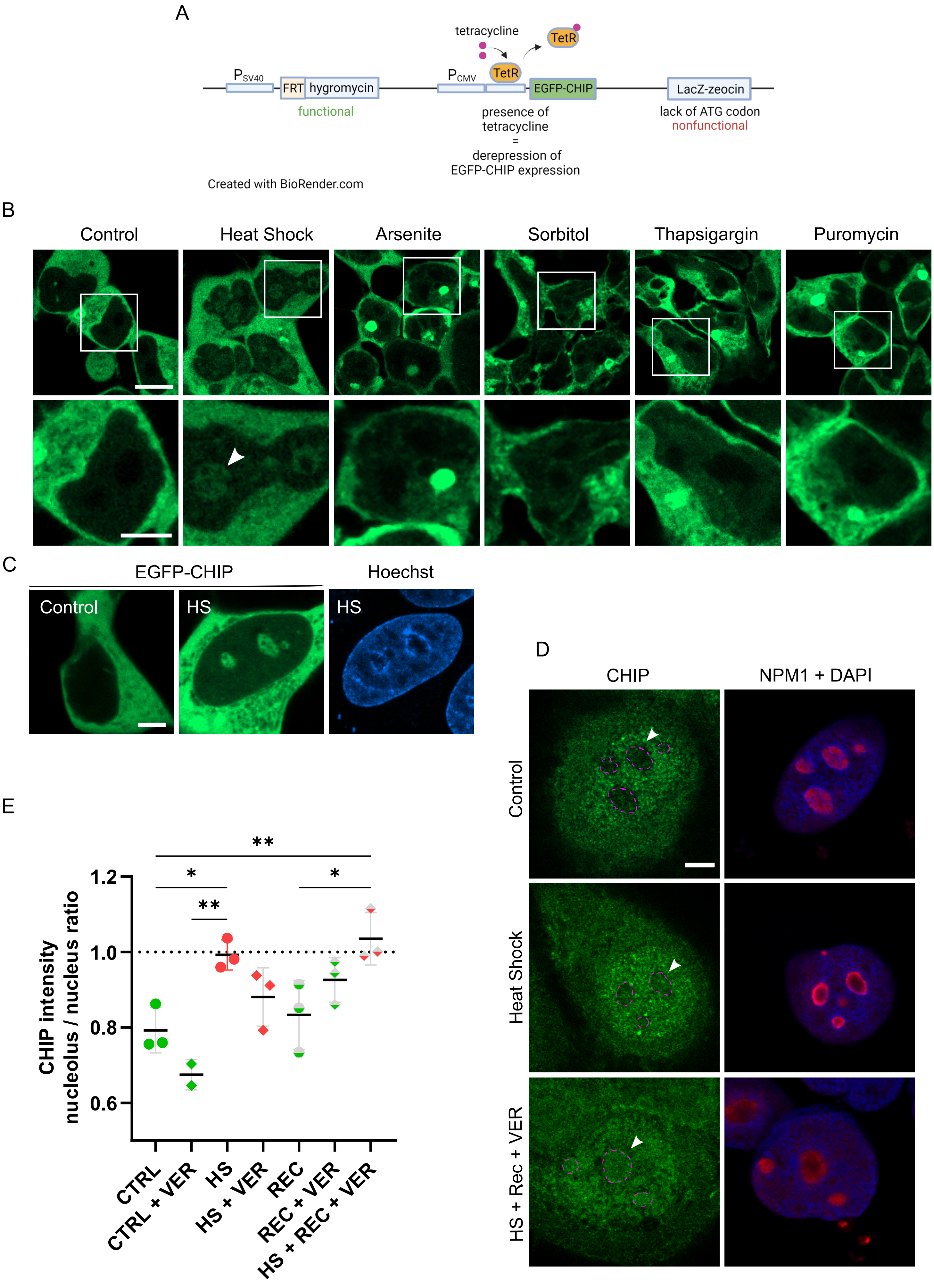

### Supplemental Figure 2

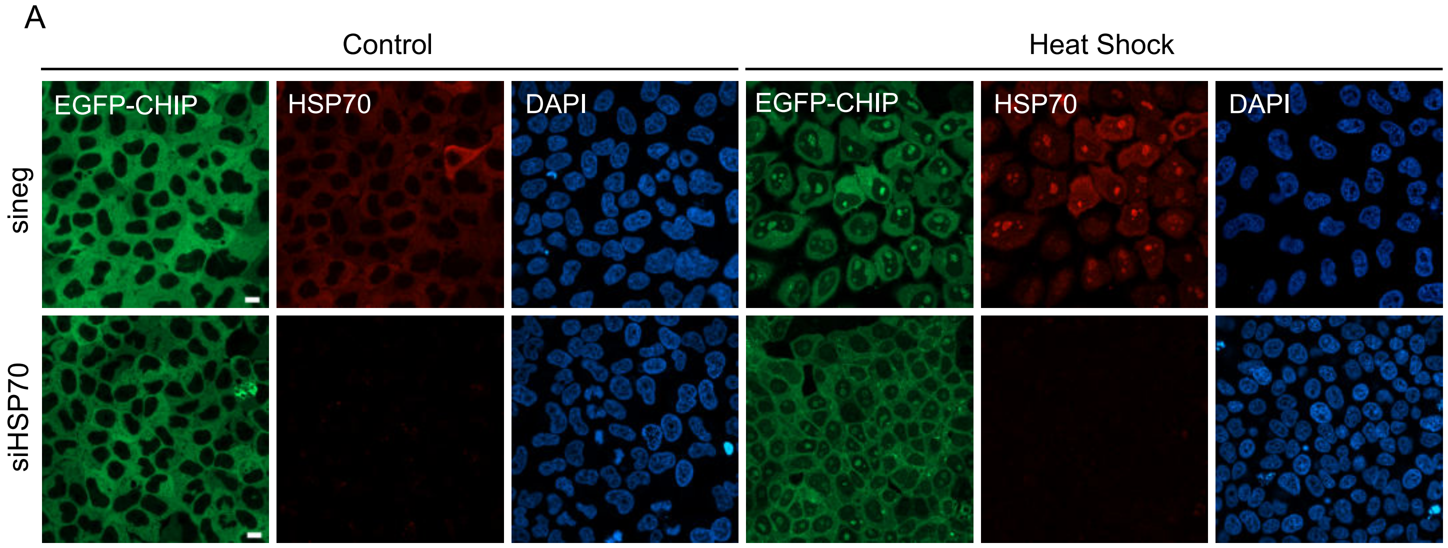

### Supplemental Figure 3

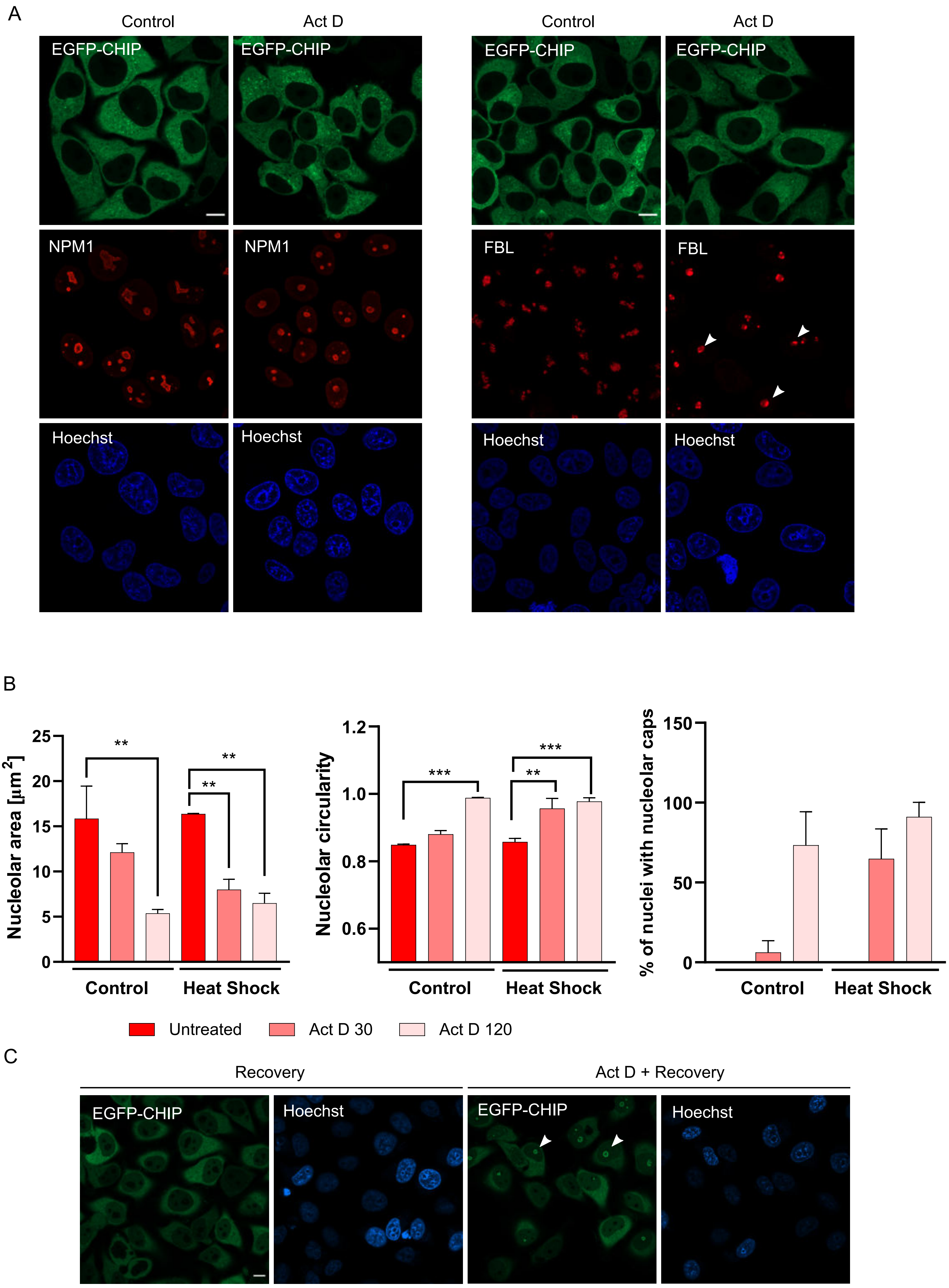

### Supplemental Figure 4

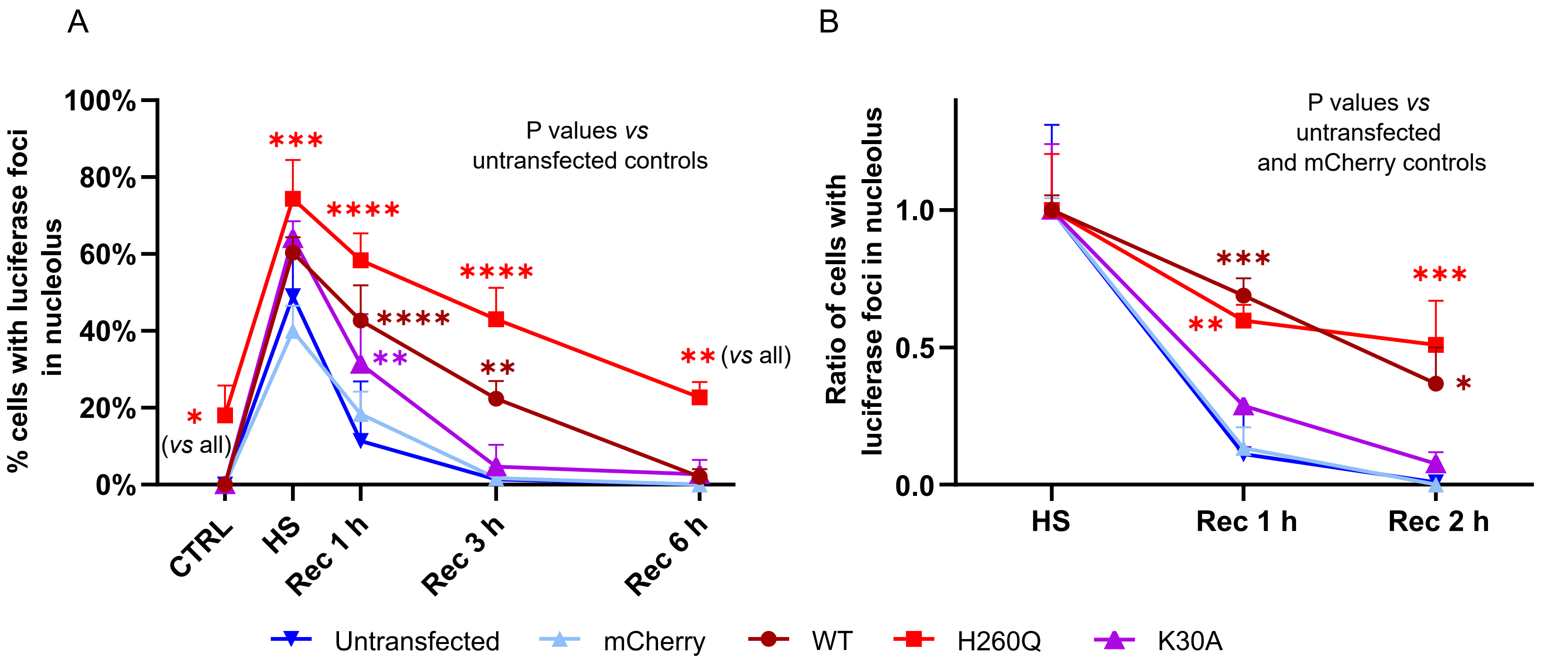

### Supplemental Figure 5

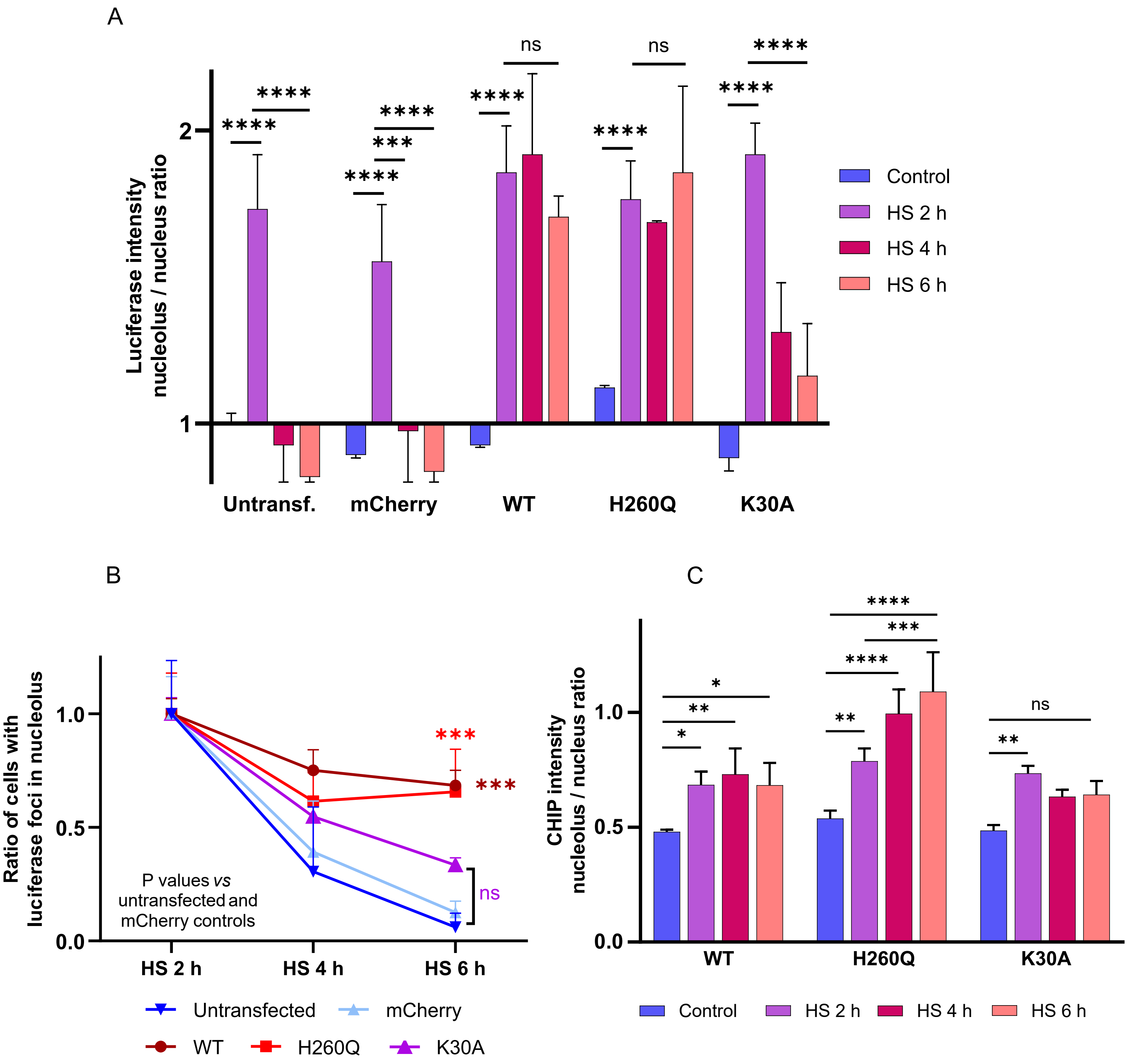
